# Whole-Body Networks: A Holistic Approach for Studying Aging

**DOI:** 10.1101/2024.07.23.604767

**Authors:** Orestis Stylianou, Johannes M. Meixner, Tilman Schlick, Colin M. Krüger

## Abstract

Aging is a multiorgan disease, yet the traditional approach is to study each organ in isolation. Such organ-specific studies allowed us to gather invaluable information regarding the pathomechanisms that contribute to senescence. But we believe that a big-picture exploration of the whole-body network (WBN) during aging could be complementary. In this study, we analyzed the functional magnetic resonance imaging (fMRI), breathing rate and heart rate time series of a young and an elderly group during eyes-open resting-state. By exploring the time-lagged coupling between the different organs we constructed WBNs. First, we showed that our analytical pipeline could identify regional differences in the networks of both populations, allowing us to proceed with the remaining of the analysis. By comparing the WBNs of young and elderly, a complex relationship emerged where some connections were stronger and some weaker in the elderly. Finally, the interconnectivity and segregation of the WBNs negatively correlated with the short-term memory of the young participants. This study: *i*) validated our methods, *ii*) identified differences between the two groups and *iii*) showed correlation with behavioral metrics. We are at the edge of a paradigm shift on how aging-related research is conducted and we believe that our methodology should be implemented in more complex mental and/or physical tasks to better demonstrate the alterations of WBNs as we age.

## Introduction

In the last few centuries, the standard of living has increased across the board, resulting in higher life expectancy. In tandem with that, we have also witnessed the proliferation of diseases that appear in the later decades of life (e.g. heart failure, tumors and diabetes mellitus type 2). For several decades these diseases were regarded as a logical byproduct of aging, a constant reminder that the rate of disorder/dysregulation in the human body always increases as we age. Yet, in the last few years scientists started calling for classifying aging as just another disease (Bulterijs et al. 2015; Zhavoronkov and Bhullar 2015). Just as in every disease, scientists try to unravel its pathomechanisms. Initially, such explorations were constrained on organ-specific dysregulations. These studies have been invaluable helping us understand and combat several age-related limitations that people face. Nonetheless, the need of a big-picture exploration of aging as a multiorgan disease seems to be more fitting, as recently shown by Tian et al. (Tian et al. 2023). In this seminal study, bidirectional influences were observed between the biological age of brain and visceral organs. It is then clear that a paradigm shift in how we study aging is imminent. This new perspective is complementary to preexisting organ-specific observations and shows great potential for future decoding of how we age and what we can do to prevent it.

The human body is a complex, interconnected system whose components are in dynamic equilibrium; meaning that changes in one compartment will result in a cascade of changes throughout the whole body. In line with that, the field of network physiology emerged (Ivanov and Bartsch 2014). Under the network physiology formalism, the human body can be represented as a whole-body network (WBN). Each WBN is a constellation of nodes interconnected via edges. Every node represents a different subsystem of the human body. These subsystems can be whole organs or, in special cases, parts of an organ. The assignment of smaller divisions to individual nodes is especially prominent in the brain, due to its regionally specialized functions. For example, a simple WBN can consist of a set of neural nodes, one cardiac node and one pulmonary node. Activity from every node is recorded in the form of signals. These time series represent the temporal evolution of the node. In our example, this would correspond to the brain, cardiac and pulmonary activity. The edges (i.e. connections) of the network are drawn by estimating the statistical interdependence of the recorded time series. Such methods have already been used to study the stress levels in the neonatal ICU (Lavanga et al. 2020) as well as the survivability of liver (Tan, Montagnese, and Mani 2020) and renal (Liu et al. 2021) failure patients. Most importantly, it was shown that transition from resting-state to task conditions could indeed be captured in the changes of WBNs (Pernice et al. 2019, 2021; Zanetti et al. 2019).

In this study, we compared the WBNs of young and elderly populations, as they were constructed based on neuronal, cardiac and pulmonary time series. Age-related alterations in the brain (Peters 2006), heart (Cheitlin 2003) and lungs (Sharma and Goodwin 2006) can be seen even in healthy individuals, hence we expected alterations to be observed in the WBNs of the studied population. We also investigated the possible correlations of WBNs’ architecture and cognitive performance – as it was recorded from an array of tests – expecting a relationship to emerge.

## Methods

### Dataset

We used the freely available LEMON Dataset (Babayan et al. 2019), which included a plethora of biosignals. For our analysis, we included only the preprocessed eyes-open resting-state fMRI time series accompanied by simultaneously recorded blood pressure and respiration. Subjects with missing blood pressure and/or respiration data were excluded from the analysis. Further, the subjects *sub-032330, sub-032374* and *sub-032489* were excluded due to bad quality respiration time series, while *sub-032355, sub-032463, sub-032481, sub-032493* and *sub-032519* were excluded due to bad quality blood pressure time series. We divided the remaining sample into two groups of *n* = 34 elderly (≥60 years old, 22 men) and *n* = 42 young (≤25 years old, 32 men) subjects. Middle-aged participants (i.e. between 25 and 60 years old) were excluded in order to increase the discriminatory power of our analysis. The subject age was given in a range (e.g. 65-70 years) hence exact ages of the two groups were not possible.

Prior to the fMRI recording the participants had to perform an array of cognitive assessment tests. One of them was the California Verbal Learning Task (CVLT) (Niemann et al. 2008), which investigated the participants’ short-term memory. The instructor read 16 words and then the subject had to recall as many words as possible from this list. This was repeated four more times, using the same list of 16 words. Using the available data, we extracted the number of correct recalls the subject had in the final (i.e. the fifth) trial. Several other tests were also available, but since they did not yield statistically significant results, we decided to include them in the **Supplementary Material** only.

We parcellated the already preprocessed (and already normalized to the MNI 152 1mm standard space) fMRI data into the seven resting-state networks (RSNs) identified by Yeo et al. (Thomas Yeo et al. 2011), using CONN (Whitfield-Gabrieli and Nieto-Castanon 2012). These RSNs were the: *i*) Default Mode Network (DMN) *ii*) Frontoparietal Network (FPN) *iii*) Limbic Network (LN) *iv*) Ventral Attention Network (VAN) *v*) Dorsal Attention Network (DAN) *vi*) Sensorimotor Network^1^ (SMN) and *vii*) Visual Network (VN). The purpose of this parcellation was to reduce the dimensionality of the data in a functionally meaningful way (van den Heuvel and Hulshoff Pol 2010). As the name suggests, DMN is the default mode of the brain, i.e. its components are active during resting-state while their activity decreases during mental tasks (Buckner, Andrews-Hanna, and Schacter 2008). FPN is involved in executive control and information integration (Vincent et al. 2008). VAN and DAN are activated when increased attention is required, with VAN being responsible for notable/unexpected triggers (Fox et al. 2006) and DAN for voluntary attention (Fox et al. 2006). LN is involved in diverse functions such as emotion, cognition and memory (Kötter 2003). SMN is responsible for voluntary movement and sensation (Kandel, Schwartz, and Jessell 2012). VN is the starting point of visual processing in the brain (Furman 2014).

We identified peaks in the blood pressure and respiration signals indicating the beginning of a new cardiac and respiratory cycle, respectively. Due to trends that did not allow for accurate local maxima estimation for the whole recording, we split the respiration time series into 60s long non-overlapping segments. In every segment we identified the local maxima of the signal using the MATLAB function *peakfinger* (Yoder 2024). As this was not the case with the blood pressure signal, we estimated the systolic peaks of the whole recording using the function *BP_annotate* (Laurin 2024). The number of peaks of blood pressure and respiration signals were counted for every 1.4s. This resulted in heart rate (HR) HR and respiration rate (RR)time series with the same sampling rate as that of the fMRI signals (**Figure 1**). Nine minutes were selected for the analysis, starting from 30s until 9m30s of the fMRI, HR and RR signals.

**Figure 1.**
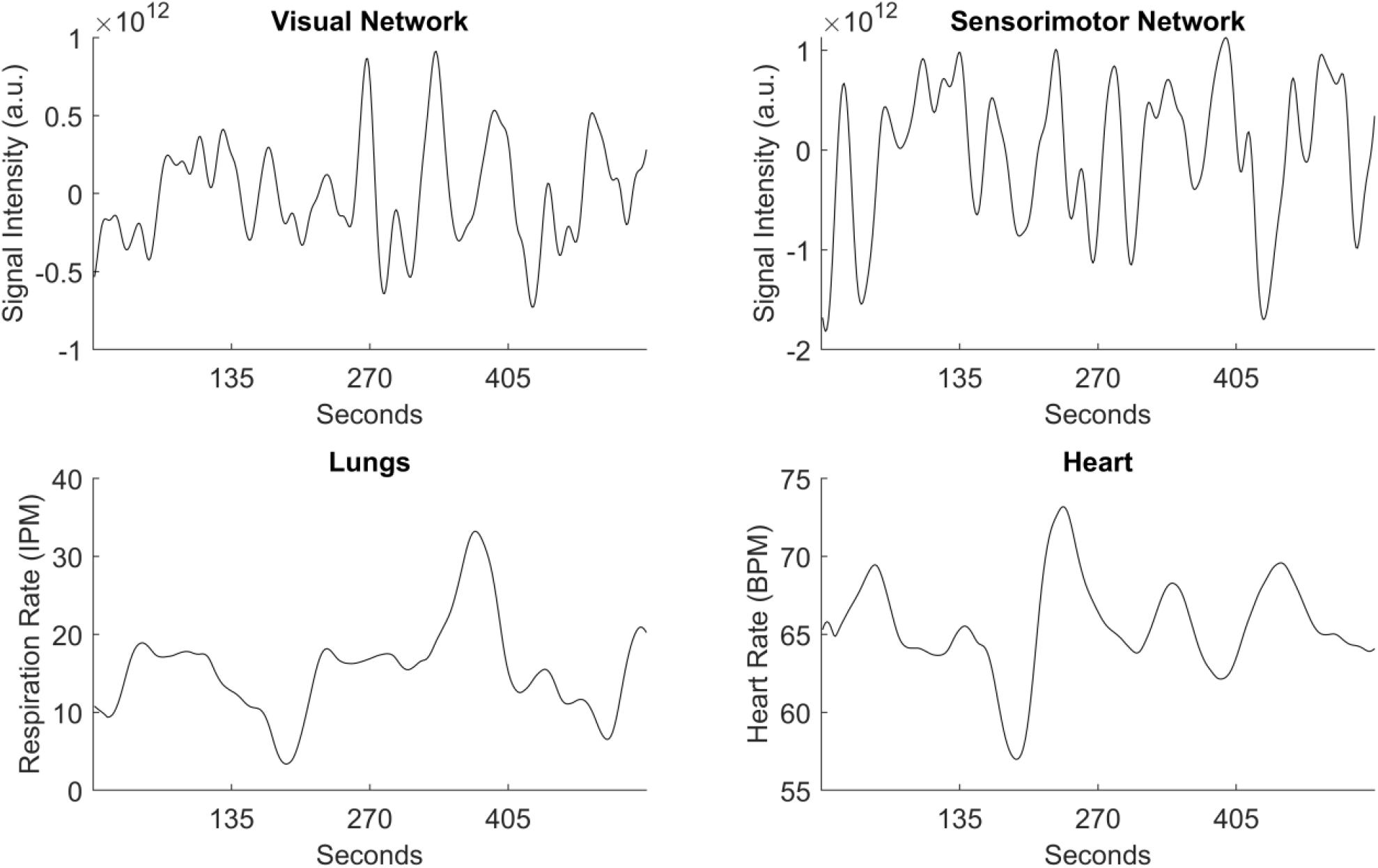
Example of the signals used in this study. The top signals correspond to fMRI activity from the visual (VN) and sensorimotor (SMN) network. The bottom signals correspond to respiration (Lungs) and heart (Heart) rate. a.u.: arbitrary unit, IPM: inspirations per minute, BPM: beats per minute

### Whole-Body Networks Construction

To construct WBNs we had to estimate the statistical interdependence of the fMRI, HR and RR signals. The simplest solution would be to use Pearson’s correlation, which is often employed in the construction of brain networks (Fornito, Zalesky, and Bullmore 2016). Brain regions are connected through rapid-firing interneurons, where activity change in one area would almost instantly correspond to activity changes in its connected regions. This is not the case when multiple organs are studied. For example, the information of emotionally disturbing imagery will first reach the visual cortex and through there it will propagate in a fast fashion to the hypothalamus that regulates the autonomic nervous system. From there, sympathetic fibers will carry the stimulus to the cardiorespiratory centers leading to increases in HR and RR. At the same time activation/deactivation of specific brain regions will be followed by alterations in the cerebral flow reflected in the fMRI signal. Due to the inherent time delays of the system, it is then clear that the zero-lag Pearson’s correlation is probably not a good estimator for WBNs. A better alternative would have been the cross-correlation, where the time lag is considered during the estimation of statistical interdependence. However, rather than a high cross-correlation value, which is maximized in greatly different lags every few moments, we believe that the stability of the connection is a more accurate estimator of strong connection between two regions.

Due to the aforementioned reasons we decided to use time delay stability (TDS), which has been specifically designed to construct WBNs (Bashan et al. 2012). We had nine signals (seven fMRI, HR and RR), each nine minutes long. Every pair of signals was segmented into 30s windows with 50% overlap (i.e. 35 segments in total). For every segment we removed the linear trends of the signal and then rendered them dimensionless by calculating the Z-scores of the time series. We then found at which lag (termed *optimal lag* in the remainder) is the absolute cross-correlation maximum between the two signals. This resulted in a 35-element vector with optimal lags, one for each window. For every window we compared the optimal lag with the following segments. If the difference between two optimal lags was smaller than 7s, the connection between the two pairs of signals was considered stable. Of course, we should not expect a stable connection to be maintained throughout the whole recording. Hence, for every segment only the following five segments, i.e. 150s, were used to indicate the stability of the connection. If in at least three out of the five segments (75%) the connection was stable (i.e. optimal lag smaller than 7s) the connection was considered truly stable. Finally, we calculated the mean of truly stable connections between segments. For example, a TDS value of 0.3 means that 30% of the segments had truly stable connections with their neighboring segments. The choice of 30s window length, 7s optimal lag and 75% percentage cutoff were selected based on an exploratory analysis. An illustration of the important steps of TDS can be seen in **Figure 2**.

**Figure 2.**
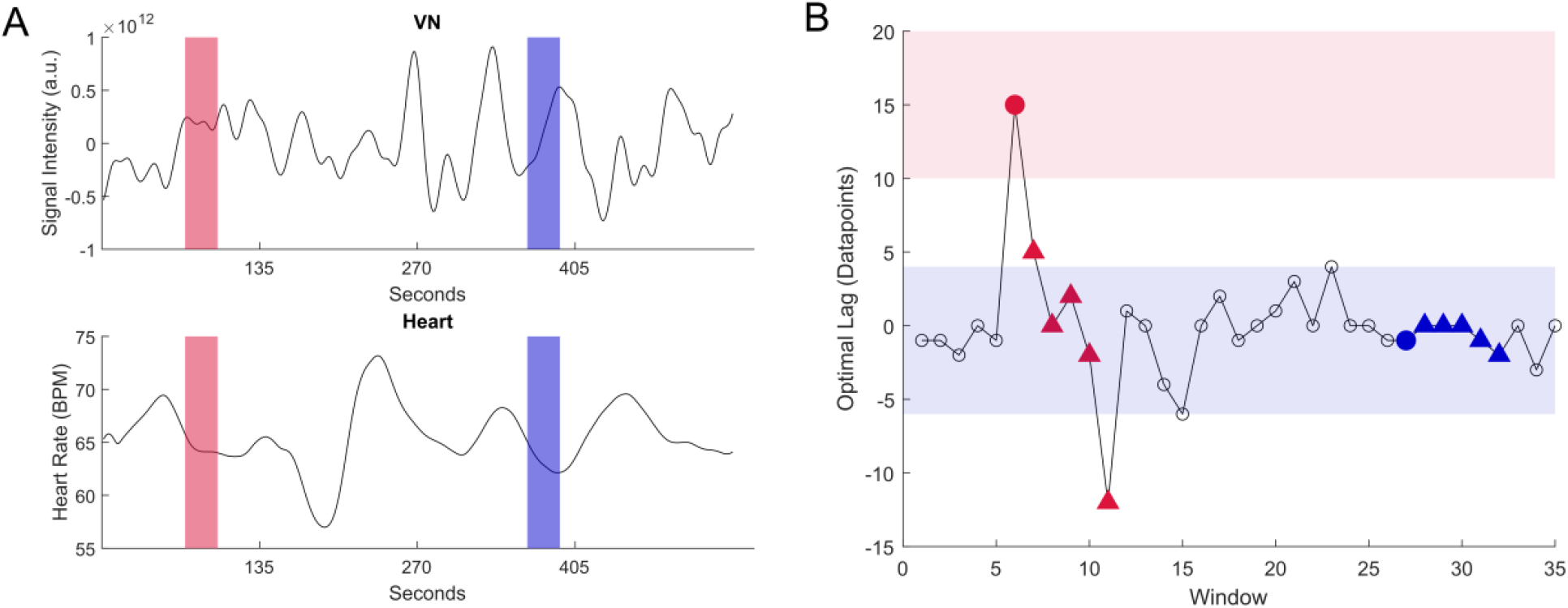
Demonstration of time delay stability (TDS). We started with the signals of visual network (VN) and heart rate (Heart), where we estimated the TDS in two different pairs of windows (red and blue) [Panel A]. In both sets of windows, we removed linear trends and normalized the time series. Then we estimated the optimal lag (where the absolute cross-correlation is maximized) and plotted it [Panel B]. The optimal lag for each window can be seen as a circle. We used the following five segments (50% overlap) (triangles) to determine if a connection is stable. A connection is stable if three out of the five triangles are within the shaded areas. The shaded areas range from (optimal lag – 5 datapoints) to (optimal lag + 5 datapoints). The 5 datapoints cutoff corresponds to 7s of lag in the time domain (7s*sampling rate ≈ 5 datapoints). In this example, we see that the connection is stable only for the blue segment. a.u.: arbitrary unit, BPM: beats per minute

Every WBN was constructed as a 9×9 matrix, representing the nine regions of the network (DMN, FPN, LN, VAN, DAN, SMN, VN, Heart and Lungs). We then calculated network metrics that can summarize the properties of the constructed WBNs (Rubinov and Sporns 2010). More specifically, we estimated the node degree (*D*), clustering coefficient (*C*) and path length (*L*). *D* indicates how interconnected one network node is, with higher values corresponding to higher interconnectivity. A similar metric is *C* which shows the number of neighboring nodes that are neighbors to each other. Higher values of *C* are found in nodes with higher segregation. Finally, *L* is the average shortest path a node has to transverse to reach all other nodes of the network. Highly integrated nodes have low *L*. Then, we estimated their average values, so for every WBN we ended up with mean node degree 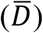, mean clustering coefficient 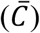 and mean path length 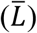.

### Statistical Evaluation

For the present purpose, we were interested only in 15 connections (seven Brain-Lungs, seven Brain-Heart and one Heart-Lungs interactions). Before comparing the connection strength (estimated by TDS) between the two age groups (e.g. DMN-Heart elderly vs DMN-Heart young), we wanted to validate that our analytical pipeline could capture regional differences in the constructed networks (e.g. DMN-Heart elderly vs VAN-Lungs elderly). This was achieved through a series of multiple comparisons of paired-sample t-tests or Wilcoxon signed rank test (depending on the normality of the distributions estimated using Lilliefors test). The estimated *p*-values were corrected using the Benjamini Hochberg (BH) correction. We then compared the connections between the two groups (e.g. DMN-Heart elderly vs DMN-Heart young) through a series of multiple comparisons of two-sample t-tests or Wilcoxon rank sum test (depending on the normality of the distributions estimated using Lilliefors test). The estimated *p*-values were corrected using the BH correction.

Before exploring any correlations with the CVLT scores, we had to make sure that the two populations had indeed different scores. We contrasted the CVLT scores of the two populations using Wilcoxon rank sum test, because the distribution of CVLT scores in the young population was not normally distributed (estimated using Lilliefors test). Finally, we wanted to see if the CVLT scores would correlate with the WBNs’ architecture in the two groups. We calculated the Spearman correlation or Pearson’s correlation between the CVLT scores and estimated network metrics (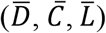) of our two populations. Spearman’s correlation was selected when at least one of the distributions in question was not normally distributed, estimated using Lilliefors test. Otherwise, we used Pearson’s correlation. For every network metric we obtained two *p*-values, one for each population. These two *p*-values were corrected using the BH correction.

## Results

Initially, we explored the regional variability captured by our analytical pipeline, i.e. how different regions of the same network vary. As shown in **Figure 3**, 15% (16 out of 105) and 17% (18 out of 105) of the connections were significantly different in the young and elderly populations, respectively. Especially prominent is the Lungs-Heart connection which is significantly higher than most of the other connections. We then investigated how regional interconnectivity can vary between the two groups, where we found that three connections were significantly different (**Figure 4**). The DAN-Heart (BH-corrected *p*-value=0.046) and DMN-Lungs (BH-corrected *p*-value=0.046) connections were higher in the young individuals, while the VAN-Lungs (BH-corrected *p*-value=0.023) connection was higher in the elderly individuals. We also see that the CVLT scores of the young group were significantly higher than that of the elderly (*p*-value<10^−5^) (**Figure 5**). Finally, significant correlations were observed only in the young group, where 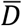 was negatively (*r*=-0.4, BH-corrected *p*-value=0.02) and 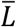 was positively (*r*=0.41, BH-corrected *p*-value=0.02) correlated with the CVLT scores (**Figure 6**). No correlations were observed in the elderly group, whose correlation results can be found in the **Supplementary Material**.

**Figure 3.**
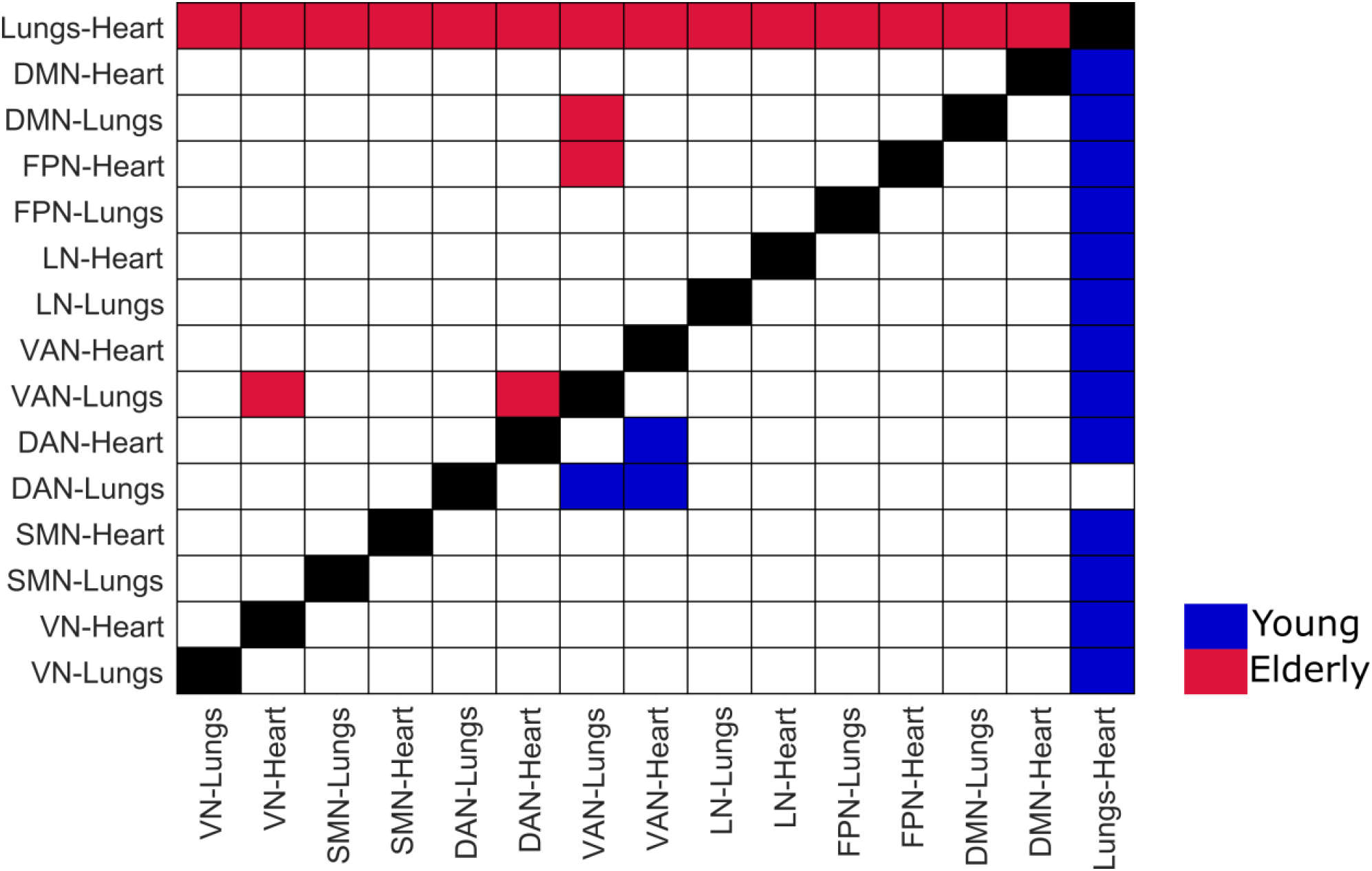
Regional variability of the constructed whole-body network. The red boxes correspond to connections that were significantly different in the elderly population. The blue boxes correspond to connections that were significantly different in the young population. White boxes represent non-significant differences. DMN: Default Mode Network, FPN: Frontoparietal Network, LN: Limbic Network, VAN: Ventral Attention Network, DAN: Dorsal Attention Network, SMN: Sensorimotor Network, VN: Visual Network

**Figure 4.**
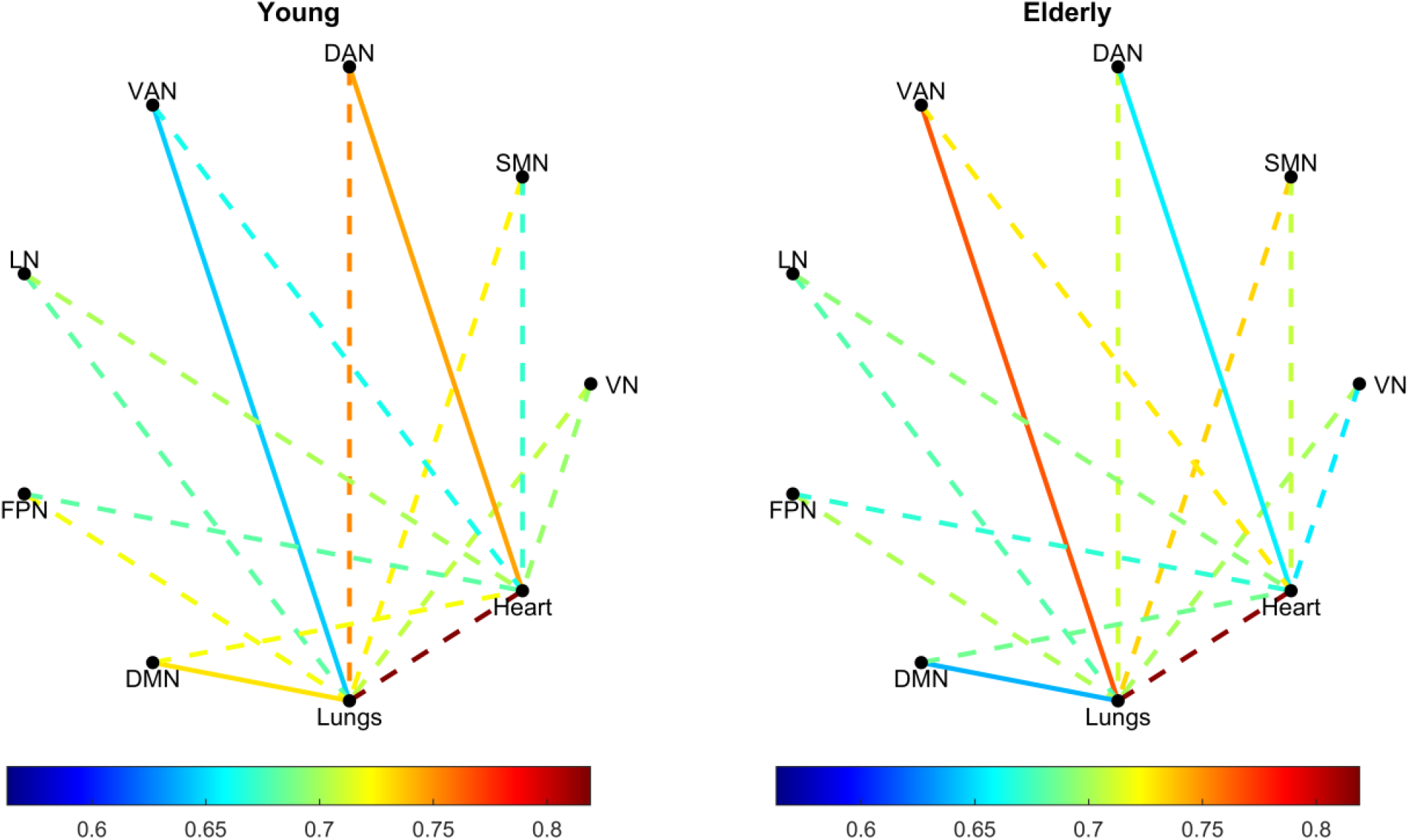
Comparison of young and elderly whole-body networks. The average time-delay stability value of the connections of the whole-body networks of the young (left) and elderly (right) is shown. Solid lines correspond to connections that were statistically different between the two populations. Dashed lines correspond to connections that were not statistically different between the two populations. DMN: Default Mode Network, FPN: Frontoparietal Network, LN: Limbic Network, VAN: Ventral Attention Network, DAN: Dorsal Attention Network, SMN: Sensorimotor Network, VN: Visual Network

**Figure 5.**
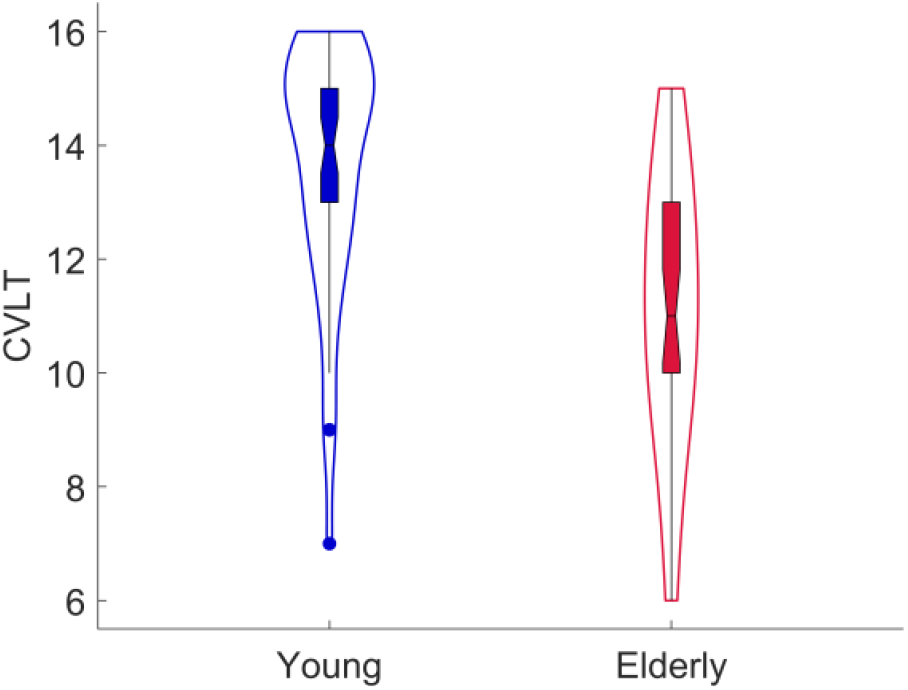
Violin plots of the California Verbal Learning Task (CVLT) scores in the young and elderly population. The instructor read 16 words and then the subject had to recall as many words as possible from this list. This was repeated four more times, using the same list of 16 words. The scores indicate the number of correct recalls the subject had in the final (i.e. the fifth) trial.

**Figure 6.**
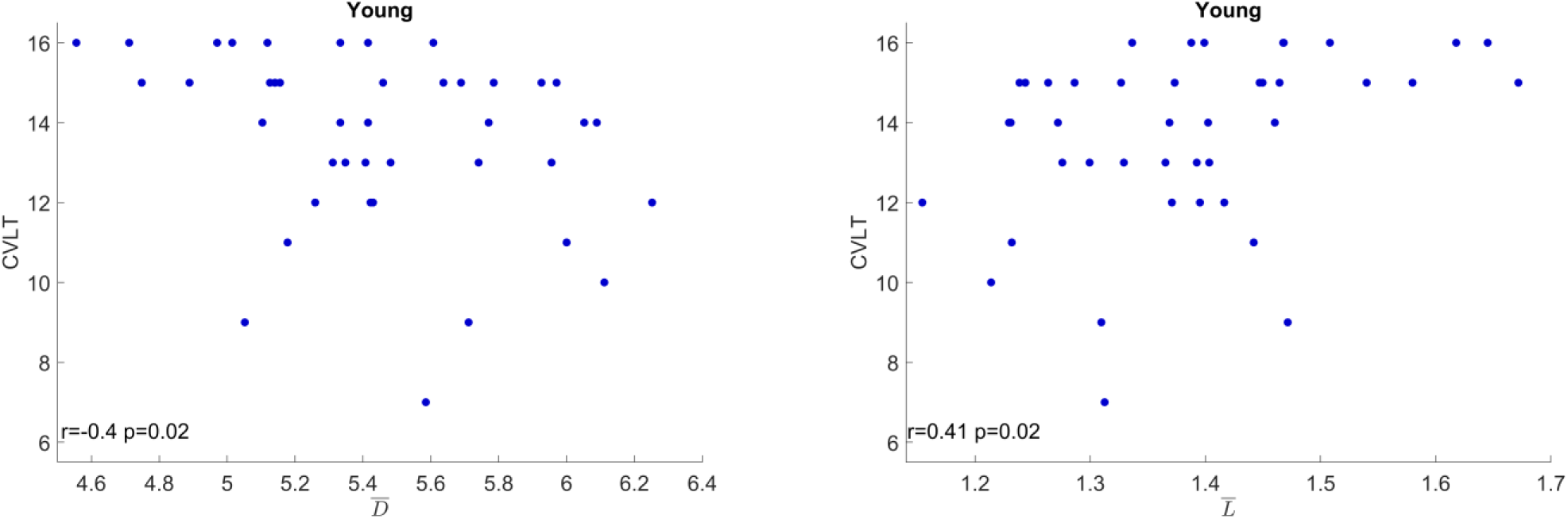
Correlations between network metrics (mean node degree or 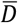 and mean path length or 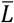) and California Verbal Learning Task (CVLT) scores in the young population.

## Discussion

Here, we explored how the new concept of WBNs can be used to study aging in a holistic manner. This was achieved by estimating the statistical interdependence between fMRI, HR and RR time series. The constructed WBNs showed both within-group and between-group differences. Finally, the interconnectivity and segregation of the WBNs negatively correlated with the short-term memory of the young participants.

A substantial number of the connections of WBNs were significantly different in both young and elderly populations (**Figure 3**). As WBNs analysis is a new field it was crucial that we validate our methodology before drawing further conclusions. The emergence of regional variability in the constructed networks warrants the exploration of this field further. Most differences were found between the Lungs-Heart and the rest of the connections. More precisely, we see that the Lungs-Heart connections are significantly higher than any other connection in both the young and elderly group. The only exception is the DAN-Lungs connection which was not statistically significant from the Lungs-Heart connection in the young group. This agrees with previous WBN studies were the Lungs-Heart connection was stronger compared to the rest during resting-state (Pernice et al. 2021; Zanetti et al. 2019). A strong interconnectivity between the lungs and heart is also in perfect alignment with our existing knowledge of the interwoven dynamics that regulate the cardiorespitory system. Here it is important to mention that TDS is not a measure of connection strength per se. As we mentioned earlier (see **Methods**), TDS indicates the stability of a connection. It is then clear that while similarities with other WBNs might be helpful, the results are heavily dependent on the used estimators of statistical interdependence. A possible variation of TDS that incorporates both stability and actual connection strength would be used to provide more information. To the best of our knowledge, this is the first study that uses WBNs to study aging. Based on the ability of our methodology to capture regional variability and the agreements with previous studies and known physiology, we believe that our approach is physiologically meaningful and appropriate to explore the WBN alterations during aging.

After comparing the strength of the connections between the WBNs of the two populations we saw that the DAN-Heart and DMN-Lungs connections were higher in the young people. On the other hand, the VAN-Lungs connection was higher in the elderly population (**Figure 4**). The decrease of connection strength during aging could indicate that the different nodes of the WBNs are decoupled. Such decoupling could be a sign of dysregulation that can be associated with the cognitive (Peters 2006), cardiac (Cheitlin 2003) and respiratory (Sharma and Goodwin 2006) alternations happening as we age. On the other hand, we also observed the increase of the VAN-Lungs connection strength. Overall, the results indicate a rather complex restructuring of WBNs, which is not just a simple decrease or increase in connectivity. During our preliminary analysis we also compared the interconnectivity 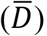, integration 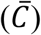 and segregation 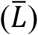 of the WBNs but no differences were found between the two groups (see **Supplementary Material**). It is believed that small-world networks (i.e. networks with high integration and high segregation) are more efficient in information transfer (Rubinov and Sporns 2010; Watts and Strogatz 1998). Hence, higher 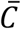 and lower 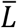 could have been expected in the young participants. A possible explanation could be that the elderly participants were healthy enough so no such alterations could be captured during rest, but they could emerge during task conditions. It is then clear that more aging studies should construct WBNs not just during rest but also during cognitive and/or physical tasks of various difficulty levels.

After confirming that the young participants performed better in the short-term memory task (**Figure 5**), we continued with our correlation analysis. We found that 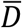 and 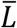 correlated negatively and positively with the CVLT scores, respectively (**Figure 6**). This was observed only in the young participants, the elderly population showed no correlations. Our results suggest that young people with lower interconnectivity and integration tend to have better short-term memory. This might seem like a counterintuitive finding, as we expect better connected systems to be more efficient. A possible explanation could be that participants with better pruning of the WBNs connections could concentrate and perform better in the task. Of course, this is only a hypothesis and due to the novelty of our investigation we cannot draw comparisons with other studies. Additionally, the recording of the biosignals was not performed during the CVLT. It is entirely possible that a different pattern could emerge if we had recordings of brain, cardiac and respiratory activity during CVLT.

Despite the clear merits of our study, some limitations should be addressed. The participants’ biosignals were recorded during resting-state. Several restrictions that the elderly population faces emerge during physical and/or mental tasks. While our current analysis helped to validate our analytical pipeline, we believe that future studies should explore age-related WBNs changes during task. Nonetheless, the parcellation of the fMRI signals to RSNs is a valid precursor to such task-related studies, since RSNs are task-specific cortical areas (van den Heuvel and Hulshoff Pol 2010; Raichle 2010). Due to TDS’s nature, our study could not estimate causal inference, which has been done in previous WBNs studies (Lin et al. 2016; Sargent et al. 2024; Zanetti et al. 2019). Another study showed that the direction of causal inference between electroencephalography (EEG) and heart rate variability (HRV) depends on the investigated EEG bands (Jurysta et al. 2003). In the future we plan to explore WBNs during task in young and elderly population using EEG, due to its high temporal resolution. It is then clear that future experiments should take causal inference into consideration. Furthermore, cardiac activity is usually estimated with HRV because it can show the interplay between the different components of the autonomic nervous system (Johnston et al. 2020). As an initial exploratory analysis we wanted to keep the analysis simple, hence we used HR. We also wanted to streamline our methodology so it can be used with wearable devices in brain computer interfaces, so minimizing analytical steps is an advantage.

## Conclusion

Here we present the findings of a multiorgan approach towards aging. Using resting-state signals recorded from the neural, cardiac and pulmonary activity we constructed WBNs. Our new methodology could capture regional differences in both the young and elderly groups. We also observed a complex pattern of alterations of the reconstructed networks in the elderly population. In the young group lower interconnectivity and integration correlated with better short-term memory. To the best of our knowledge, this is the first study exploring aging using WBNs and we believe it paves the way for further research.

## Supporting information

Correlations_clustering_coefficient.pdf

Correlations_node_degree.pdf

Correlations_path_length.pdf

Whole-Body Networks: A Holistic Approach for Studying Aging

## Author Contributions

O.S. performed data analysis and interpretation and wrote the first draft of the manuscript. J.M., T.S. and C.M.K contributed to data interpretation. C.M.K. provided conceptual guidance, supervision and funding throughout the study. All authors contributed to reviewing the manuscript and approved its final version

## Data Availability

An open dataset was used for the analysis. More information can be found out (Babayan et al. 2019).

## Footnotes

1 Despite called somatomotor by Yeo et al. this network includes also the sensory cortex, hence we modified its name.

## References

Babayan, Anahit, Miray Erbey, Deniz Kumral, Janis D. Reinelt, Andrea M. F. Reiter, Josefin Röbbig, H. Lina Schaare, Marie Uhlig, Alfred Anwander, Pierre-Louis Bazin, Annette Horstmann, Leonie Lampe, Vadim V. Nikulin, Hadas Okon-Singer, Sven Preusser, André Pampel, Christiane S. Rohr, Julia Sacher, Angelika Thöne-Otto, Sabrina Trapp, Till Nierhaus, Denise Altmann, Katrin Arelin, Maria Blöchl, Edith Bongartz, Patric Breig, Elena Cesnaite, Sufang Chen, Roberto Cozatl, Saskia Czerwonatis, Gabriele Dambrauskaite, Maria Dreyer, Jessica Enders, Melina Engelhardt, Marie Michele Fischer, Norman Forschack, Johannes Golchert, Laura Golz, C. Alexandrina Guran, Susanna Hedrich, Nicole Hentschel, Daria I. Hoffmann, Julia M. Huntenburg, Rebecca Jost, Anna Kosatschek, Stella Kunzendorf, Hannah Lammers, Mark E. Lauckner, Keyvan Mahjoory, Ahmad S. Kanaan, Natacha Mendes, Ramona Menger, Enzo Morino, Karina Näthe, Jennifer Neubauer, Handan Noyan, Sabine Oligschläger, Patricia Panczyszyn-Trzewik, Dorothee Poehlchen, Nadine Putzke, Sabrina Roski, Marie-Catherine Schaller, Anja Schieferbein, Benito Schlaak, Robert Schmidt, Krzysztof J. Gorgolewski, Hanna Maria Schmidt, Anne Schrimpf, Sylvia Stasch, Maria Voss, Annett Wiedemann, Daniel S. Margulies, Michael Gaebler, and Arno Villringer. 2019. “A Mind-Brain-Body Dataset of MRI, EEG, Cognition, Emotion, and Peripheral Physiology in Young and Old Adults.” Scientific Data 6(1):180308. doi: 10.1038/sdata.2018.308.

Bashan, Amir, Ronny P. Bartsch, Jan W. Kantelhardt, Shlomo Havlin, and Plamen Ch Ivanov. 2012. “Network Physiology Reveals Relations between Network Topology and Physiological Function.” Nature Communications 3(1):702. doi: 10.1038/ncomms1705.

Buckner, Randy L., Jessica R. Andrews-Hanna, and Daniel L. Schacter. 2008. “The Brain’s Default Network.” Annals of the New York Academy of Sciences 1124(1):1–38. doi: 10.1196/annals.1440.011.

Bulterijs, Sven, Raphaella S. Hull, Victor C. E. Björk, and Avi G. Roy. 2015. “It Is Time to Classify Biological Aging as a Disease.” Frontiers in Genetics 6. doi: 10.3389/fgene.2015.00205.

Cheitlin, Melvin D. 2003. “Cardiovascular Physiology-Changes with Aging.” The American Journal of Geriatric Cardiology 12(1):9–13. doi: 10.1111/j.1076-7460.2003.01751.x.

Fornito, Alex, Andrew Zalesky, and Edward T. Bullmore, eds. 2016. “Chapter 1 - An Introduction to Brain Networks.” Pp. 1–35 in Fundamentals of Brain Network Analysis. San Diego: Academic Press.

Fox, Michael D., Maurizio Corbetta, Abraham Z. Snyder, Justin L. Vincent, and Marcus E. Raichle. 2006. “Spontaneous Neuronal Activity Distinguishes Human Dorsal and Ventral Attention Systems.” Proceedings of the National Academy of Sciences 103(26):10046–51. doi: 10.1073/pnas.0604187103.

Furman, Moran. 2014. “Chapter 19 - Visual Network.” Pp. 247–59 in Neuronal Networks in Brain Function, CNS Disorders, and Therapeutics, edited by C. L. Faingold and H. Blumenfeld. San Diego: Academic Press.

van den Heuvel, Martijn P., and Hilleke E. Hulshoff Pol. 2010. “Exploring the Brain Network: A Review on Resting-State fMRI Functional Connectivity.” European Neuropsychopharmacology : The Journal of the European College of Neuropsychopharmacology 20(8):519–34. doi: 10.1016/j.euroneuro.2010.03.008.

Ivanov, Plamen Ch., and Ronny P. Bartsch. 2014. “Network Physiology: Mapping Interactions Between Networks of Physiologic Networks.” Pp. 203–22 in.

Johnston, Brian W., Richard Barrett-Jolley, Anton Krige, and Ingeborg D. Welters. 2020. “Heart Rate Variability: Measurement and Emerging Use in Critical Care Medicine.” Journal of the Intensive Care Society 21(2):148–57. doi: 10.1177/1751143719853744.

Jurysta, F., P. van de Borne, P. F. Migeotte, M. Dumont, J. P. Lanquart, J. P. Degaute, and P. Linkowski. 2003. “A Study of the Dynamic Interactions between Sleep EEG and Heart Rate Variability in Healthy Young Men.” Clinical Neurophysiology 114(11):2146–55. doi: 10.1016/S1388-2457(03)00215-3.

Kandel, E. R., James H. Schwartz, and T. M. Jessell. 2012. Principles of Neural Science. McGraw-Hill Professional Pub.

Kötter, Rolf. 2003. “Limbic System.” Pp. 796–99 in Encyclopedia of the Neurological Sciences, edited by M. J. Aminoff and R. B. Daroff. New York: Academic Press.

Laurin, Alexandre. 2024. “BP_annotate.” Retrieved April 4, 2024 (https://www.mathworks.com/matlabcentral/fileexchange/60172-bp_annotate).

Lavanga, Mario, Bieke Bollen, Katrien Jansen, Els Ortibus, Gunnar Naulaers, Sabine Van Huffel, and Alexander Caicedo. 2020. “A Bradycardia-Based Stress Calculator for the Neonatal Intensive Care Unit: A Multisystem Approach.” Frontiers in Physiology 11. doi: 10.3389/fphys.2020.00741.

Lin, Aijing, Kang K. L. Liu, Ronny P. Bartsch, and Plamen Ch. Ivanov. 2016. “Delay-Correlation Landscape Reveals Characteristic Time Delays of Brain Rhythms and Heart Interactions.” Philosophical Transactions of the Royal Society A: Mathematical, Physical and Engineering Sciences 374(2067):20150182. doi: 10.1098/rsta.2015.0182.

Liu, Mingxin, Véronique Legault, Tamàs Fülöp, Anne-Marie Côté, Dominique Gravel, F. Guillaume Blanchet, Diana L. Leung, Sylvia Juhong Lee, Yuichi Nakazato, and Alan A. Cohen. 2021. “Prediction of Mortality in Hemodialysis Patients Using Moving Multivariate Distance.” Frontiers in Physiology 12. doi: 10.3389/fphys.2021.612494.

Niemann, H., W. Sturm, Angelika I. T. Thöne-Otto, and Klaus Willmes. 2008. “CVLT California Verbal Learning Test. German Adaptation. Manual.”

Pernice, Riccardo, Yuri Antonacci, Matteo Zanetti, Alessandro Busacca, Daniele Marinazzo, Luca Faes, and Giandomenico Nollo. 2021. “Multivariate Correlation Measures Reveal Structure and Strength of Brain–Body Physiological Networks at Rest and During Mental Stress.” Frontiers in Neuroscience 14. doi: 10.3389/fnins.2020.602584.

Pernice, Riccardo, Matteo Zanetti, Giandomenico Nollo, Mariolin De Cecco, Alessandro Busacca, and Luca Faes. 2019. “Mutual Information Analysis of Brain-Body Interactions during Different Levels of Mental Stress.” Pp. 6176–79 in 2019 41st Annual International Conference of the IEEE Engineering in Medicine and Biology Society (EMBC).

Peters, R. 2006. “Ageing and the Brain.” Postgraduate Medical Journal 82(964):84–88. doi: 10.1136/pgmj.2005.036665.

Raichle, Marcus E. 2010. “Two Views of Brain Function.” Trends in Cognitive Sciences 14(4):180–90. doi: 10.1016/j.tics.2010.01.008.

Rubinov, Mikail, and Olaf Sporns. 2010. “Complex Network Measures of Brain Connectivity: Uses and Interpretations.” NeuroImage 52(3):1059–69. doi: 10.1016/j.neuroimage.2009.10.003.

Sargent, Kaia S., Emily L. Martinez, Alexandra C. Reed, Anika Guha, Morgan E. Bartholomew, Caroline K. Diehl, Christine S. Chang, Sarah Salama, Tzvetan Popov, Julian F. Thayer, Gregory A. Miller, and Cindy M. Yee. 2024. “Oscillatory Coupling Between Neural and Cardiac Rhythms.” Psychological Science 09567976241235932. doi: 10.1177/09567976241235932.

Sharma, Gulshan, and James Goodwin. 2006. “Effect of Aging on Respiratory System Physiology and Immunology.” Clinical Interventions in Aging 1(3):253–60. doi: 10.2147/ciia.2006.1.3.253.

Tan, Yen Yi, Sara Montagnese, and Ali R. Mani. 2020. “Organ System Network Disruption Is Associated With Poor Prognosis in Patients With Chronic Liver Failure.” Frontiers in Physiology 11(August):1–14. doi: 10.3389/fphys.2020.00983.

Thomas Yeo, B. T., Fenna M. Krienen, Jorge Sepulcre, Mert R. Sabuncu, Danial Lashkari, Marisa Hollinshead, Joshua L. Roffman, Jordan W. Smoller, Lilla Zöllei, Jonathan R. Polimeni, Bruce Fischl, Hesheng Liu, and Randy L. Buckner. 2011. “The Organization of the Human Cerebral Cortex Estimated by Intrinsic Functional Connectivity.” Journal of Neurophysiology 106(3):1125–65. doi: 10.1152/jn.00338.2011.

Tian, Ye Ella, Vanessa Cropley, Andrea B. Maier, Nicola T. Lautenschlager, Michael Breakspear, and Andrew Zalesky. 2023. “Heterogeneous Aging across Multiple Organ Systems and Prediction of Chronic Disease and Mortality.” Nature Medicine 29(5):1221–31. doi: 10.1038/s41591-023-02296-6.

Vincent, Justin L., Itamar Kahn, Abraham Z. Snyder, Marcus E. Raichle, and Randy L. Buckner. 2008. “Evidence for a Frontoparietal Control System Revealed by Intrinsic Functional Connectivity.” Journal of Neurophysiology 100(6):3328–42. doi: 10.1152/jn.90355.2008.

Watts, Duncan J., and Steven H. Strogatz. 1998. “Collective Dynamics of ‘Small-World’ Networks.” Nature 393(6684):440–42. doi: 10.1038/30918.

Whitfield-Gabrieli, Susan, and Alfonso Nieto-Castanon. 2012. “Conn: A Functional Connectivity Toolbox for Correlated and Anticorrelated Brain Networks.” Brain Connectivity 2(3):125–41. doi: 10.1089/brain.2012.0073.

Yoder, Nathanael. 2024. “Peakfinder(X0, Sel, Thresh, Extrema, includeEndpoints, Interpolate).” Retrieved April 4, 2024 (https://www.mathworks.com/matlabcentral/fileexchange/25500-peakfinder-x0-sel-thresh-extrema-includeendpoints-interpolate).

Zanetti, Matteo, Luca Faes, Giandomenico Nollo, Mariolino De Cecco, Riccardo Pernice, Luca Maule, Marco Pertile, and Alberto Fornaser. 2019. “Information Dynamics of the Brain, Cardiovascular and Respiratory Network during Different Levels of Mental Stress.” Entropy 21(3):275. doi: 10.3390/e21030275.

Zhavoronkov, Alex, and Bhupinder Bhullar. 2015. “Classifying Aging as a Disease in the Context of ICD-11.” Frontiers in Genetics 6. doi: 10.3389/fgene.2015.00326.

