## Supplementary material for "Whole-Body Networks: A Holistic Approach for Studying Aging": Correlations_clustering_coefficient.pdf

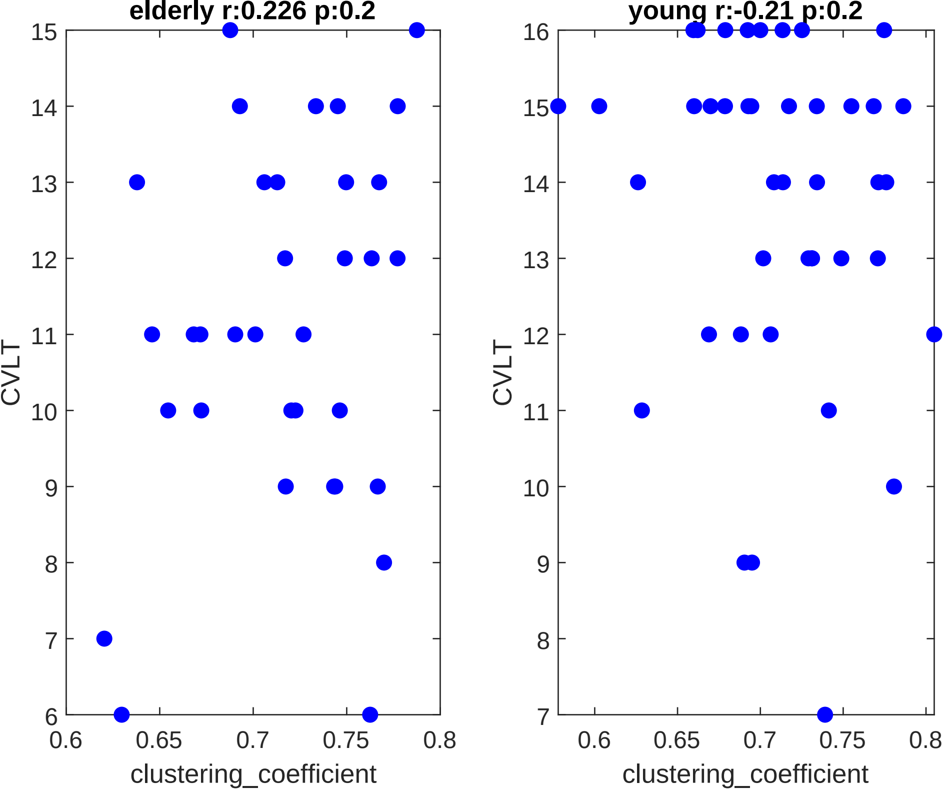

elderly r:0.226 p:0.2
young r:-0.21 p:0.2
CVLT
clustering_coefficient
15
14
13
12
11
10
9
8
7
6
16
0.6
0.65
0.7
0.75
0.8

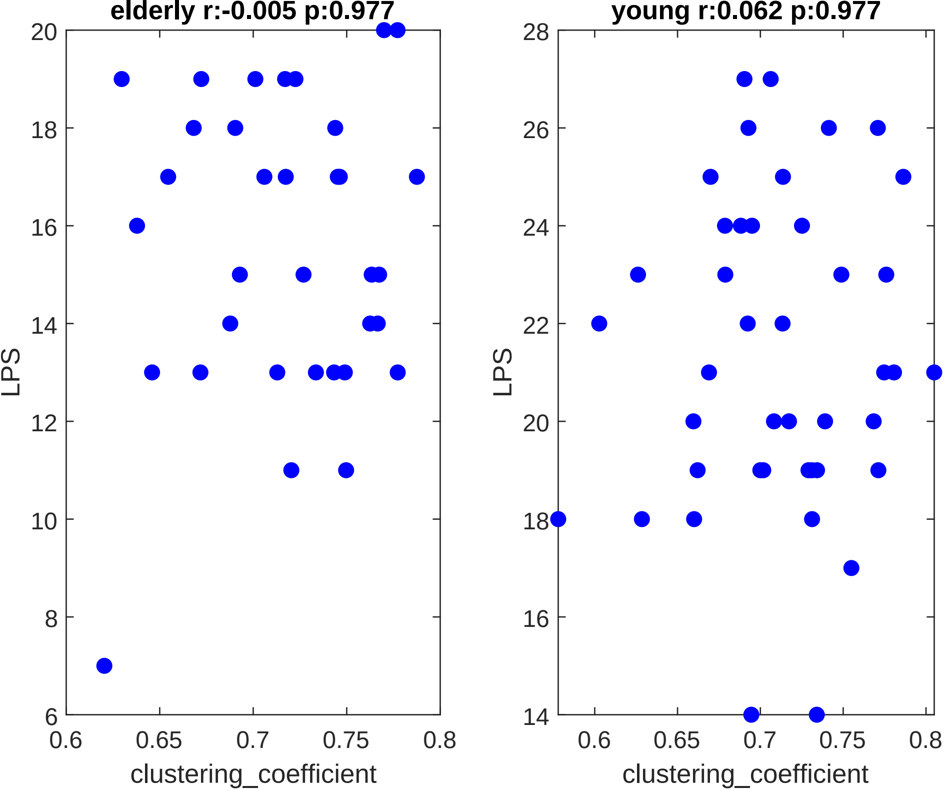

elderly r:-0.005 p:0.977
young r:0.062 p:0.977
LPS
clustering_coefficient
20
18
16
14
12
10
8
6
0.6
0.65
0.7
0.75
0.8
28
26
24
22
20
18
16
14
0.6
0.65
0.7
0.75
0.8
LPS
clustering_coefficient

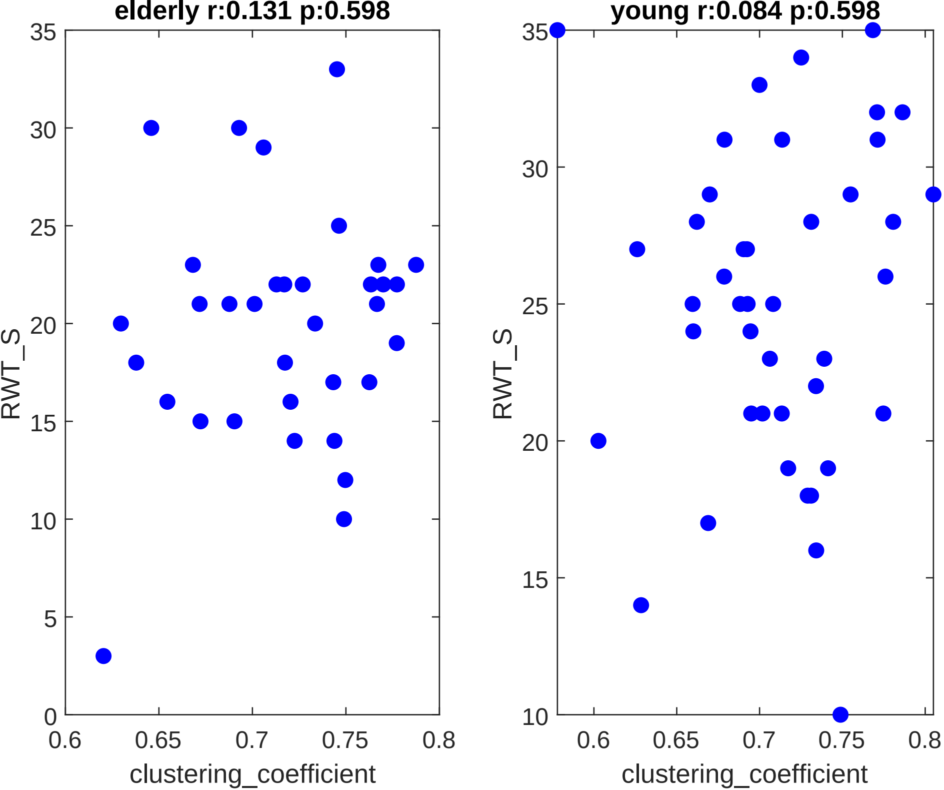

elderly r:0.131 p:0.598
young r:0.084 p:0.598
RWT_S
RWT_S
clustering_coefficient
clustering_coefficient
0
5
10
15
20
25
30
35
0.6
0.65
0.7
0.75
0.8

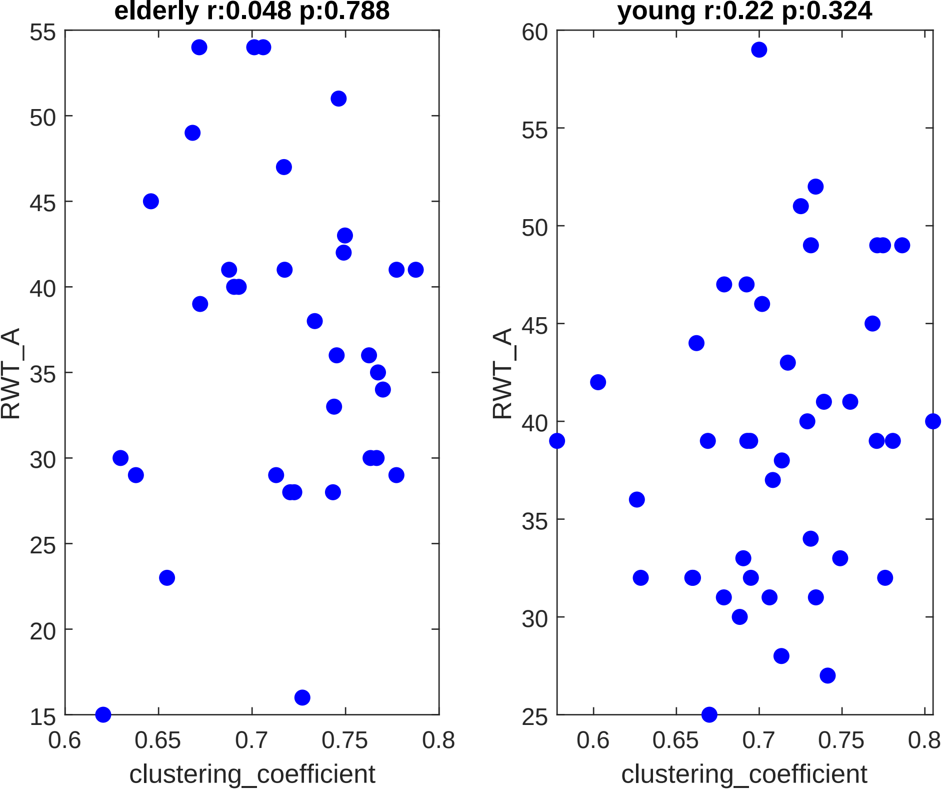
elderly r:0.048 p:0.788
young r:0.22 p:0.324
RWT_A
clustering_coefficient
55
50
45
40
35
30
25
20
15
60
0.6
0.65
0.7
0.75
0.8

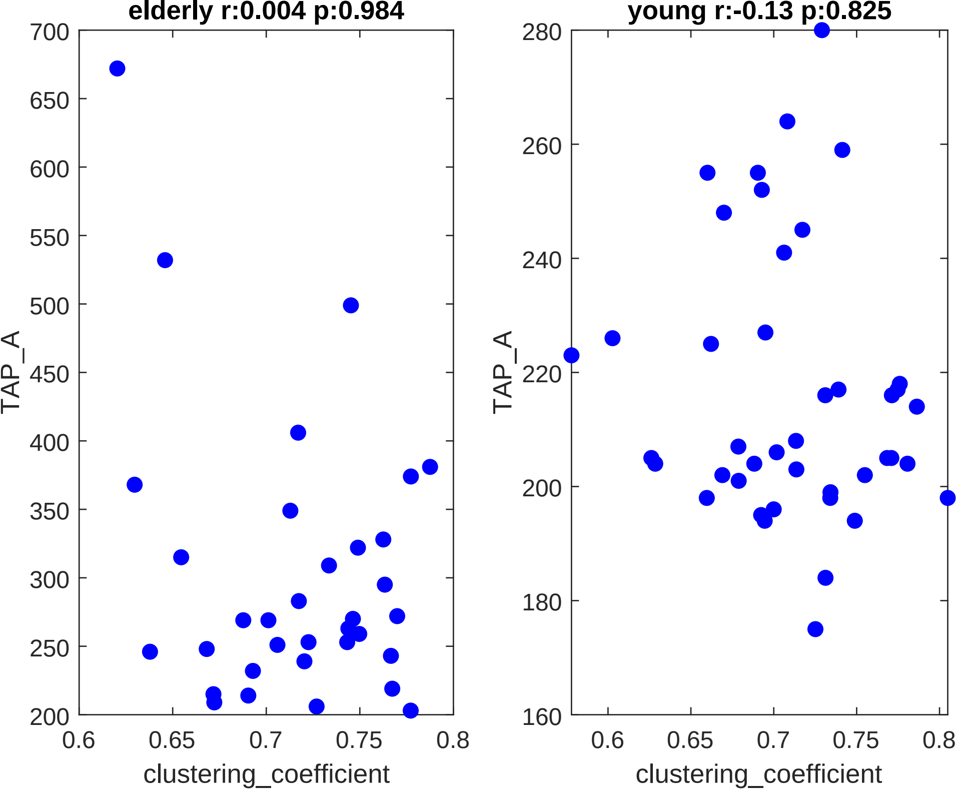

elderly r:0.004 p:0.984
young r:-0.13 p:0.825
TAP_A
TAP_A
700
650
600
550
500
450
400
350
300
250
200
280
260
240
220
200
180
160
0.6
0.65
0.7
0.75
0.8
clustering_coefficient
clustering_coefficient

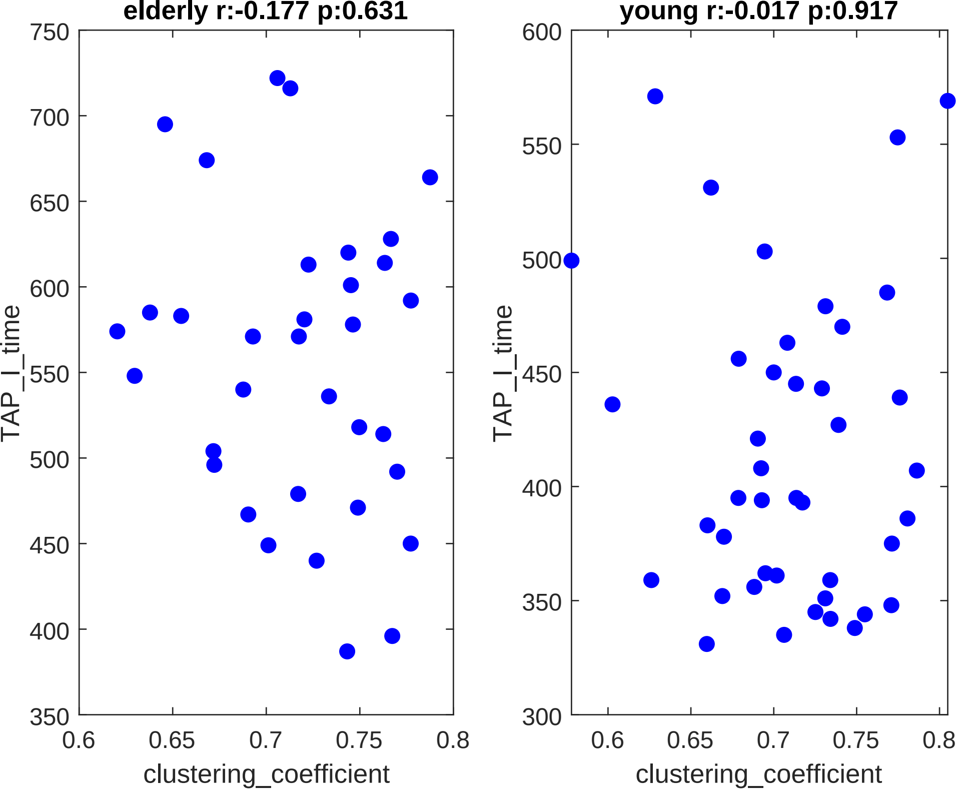
elderly r:-0.177 p:0.631
young r:-0.017 p:0.917
TAP_I_time
clustering_coefficient
750
700
650
600
550
500
450
400
350
300
0.6
0.65
0.7
0.75
0.8

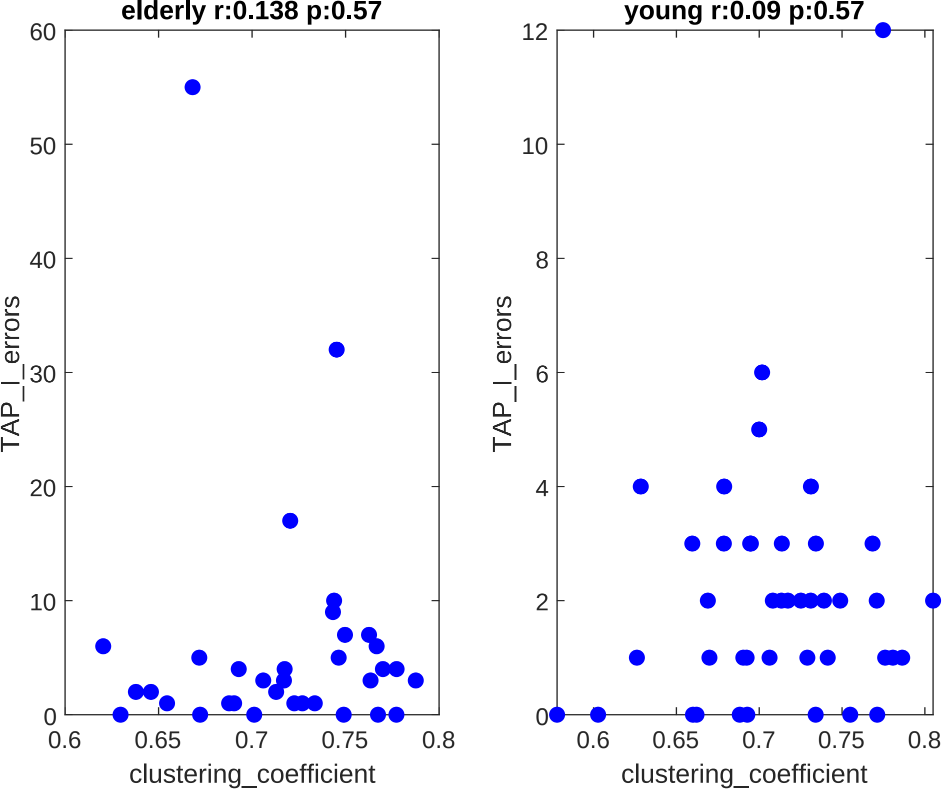

elderly r:0.138 p:0.57
TAP_I_errors
clustering_coefficient
0
10
20
30
40
50
60
0.6
0.65
0.7
0.75
0.8
young r:0.09 p:0.57
TAP_I_errors
clustering_coefficient
0
2
4
6
8
10
12
0.6
0.65
0.7
0.75
0.8

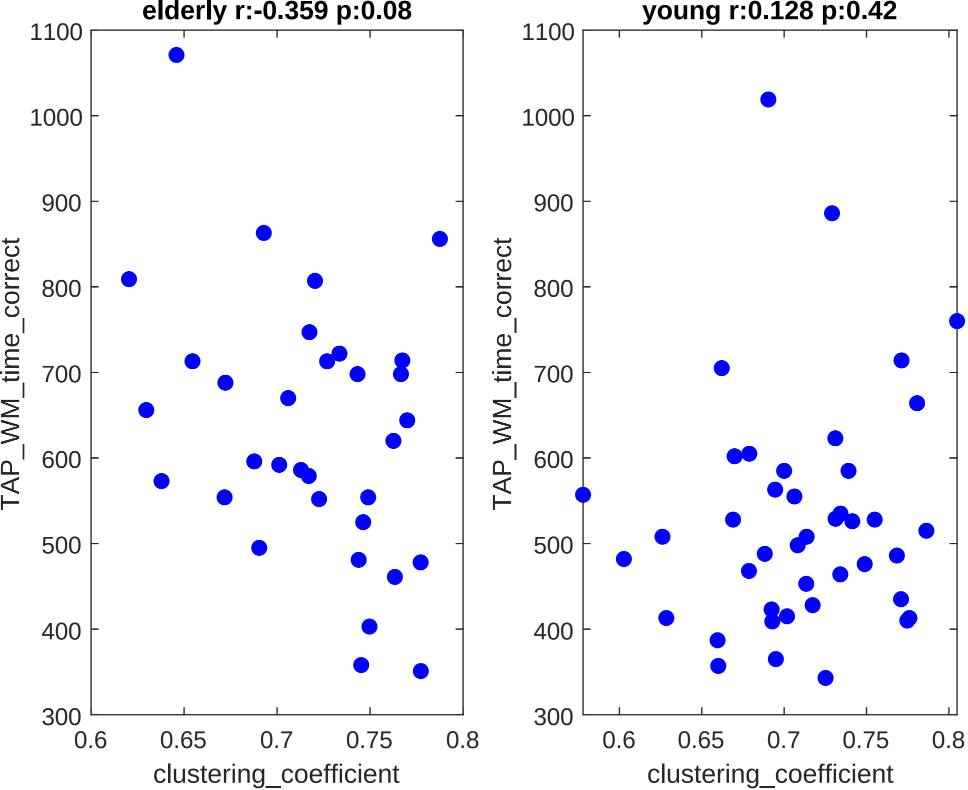
elderly r:-0.359 p:0.08
young r:0.128 p:0.42
TAP_WM_time_correct
clustering_coefficient
1100
1000
900
800
700
600
500
400
300
0.6
0.65
0.7
0.75
0.8

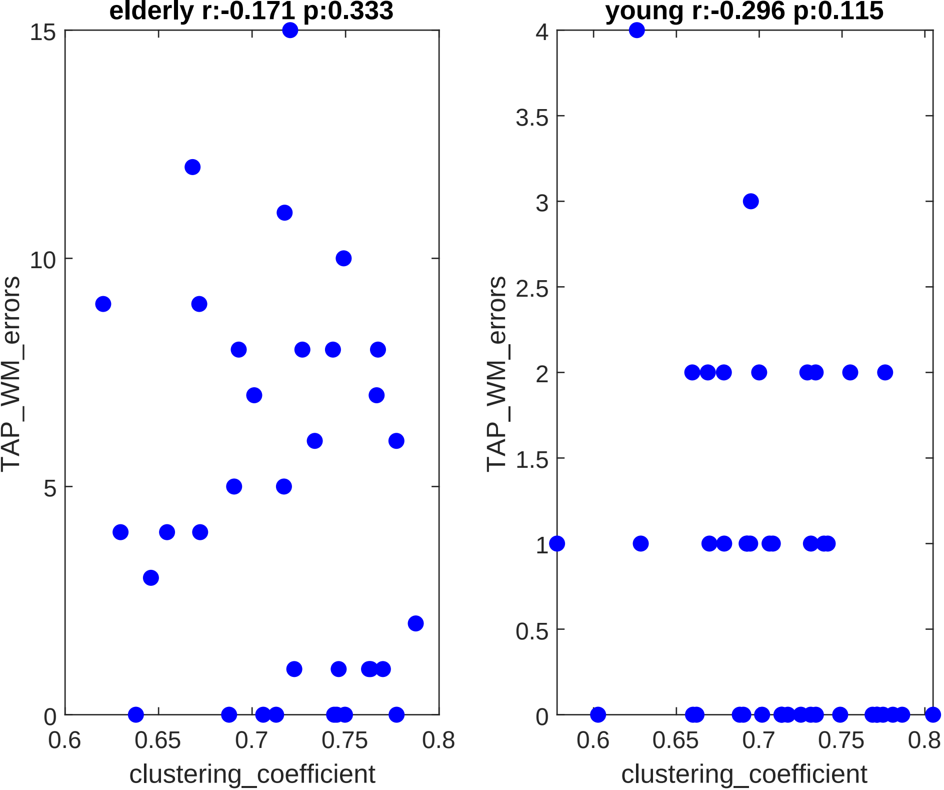

elderly r:-0.171 p:0.333
young r:-0.296 p:0.115
TAP_WM_errors
clustering_coefficient
15
10
5
0
4
3.5
3
2.5
2
1.5
1
0.5
0.6
0.65
0.7
0.75
0.8

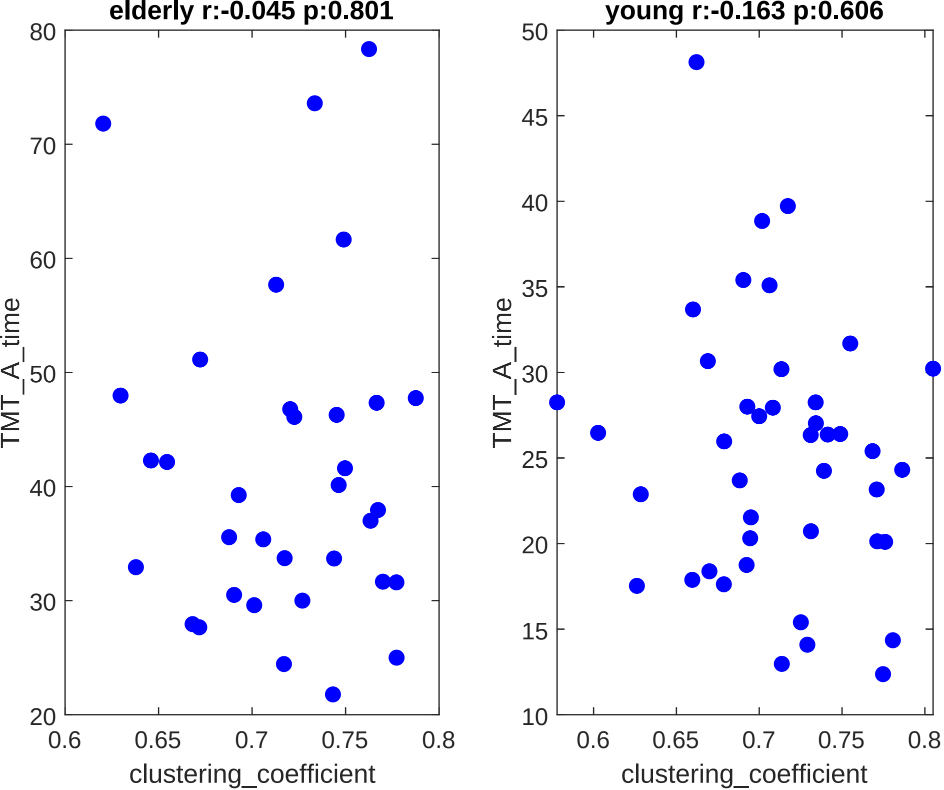
elderly r:-0.045 p:0.801
young r:-0.163 p:0.606
TMT_A_time
clustering_coefficient
80
70
60
50
40
30
20
0.6
0.65
0.7
0.75
0.8
45
35
25
15
10

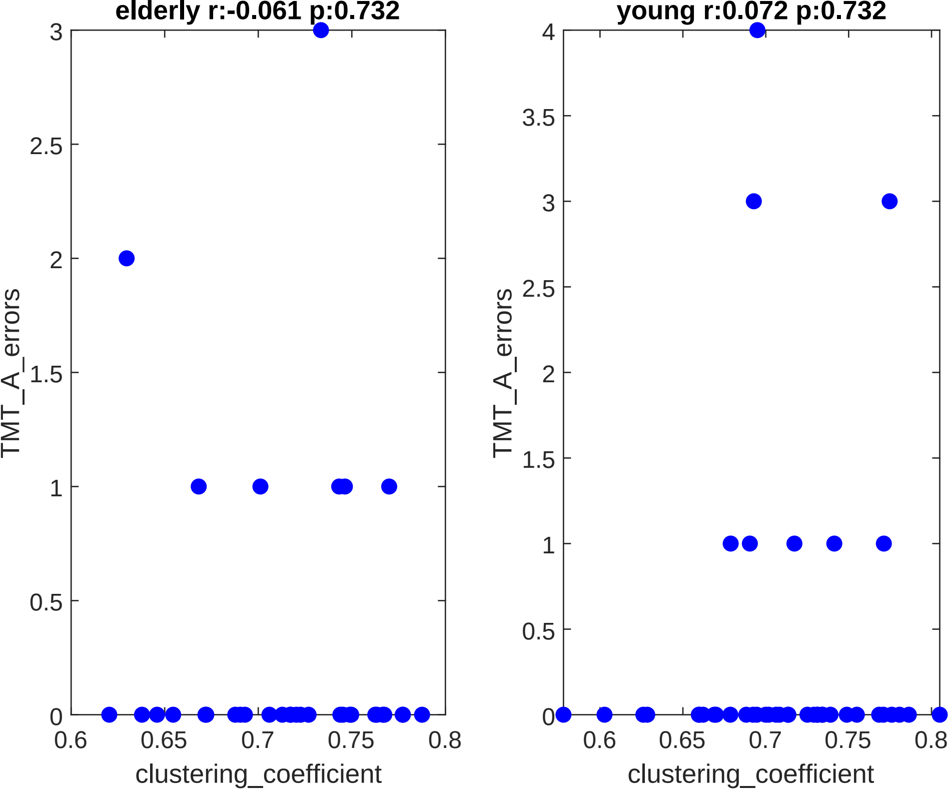

elderly r:-0.061 p:0.732
young r:0.072 p:0.732
TMT_A_errors
clustering_coefficient

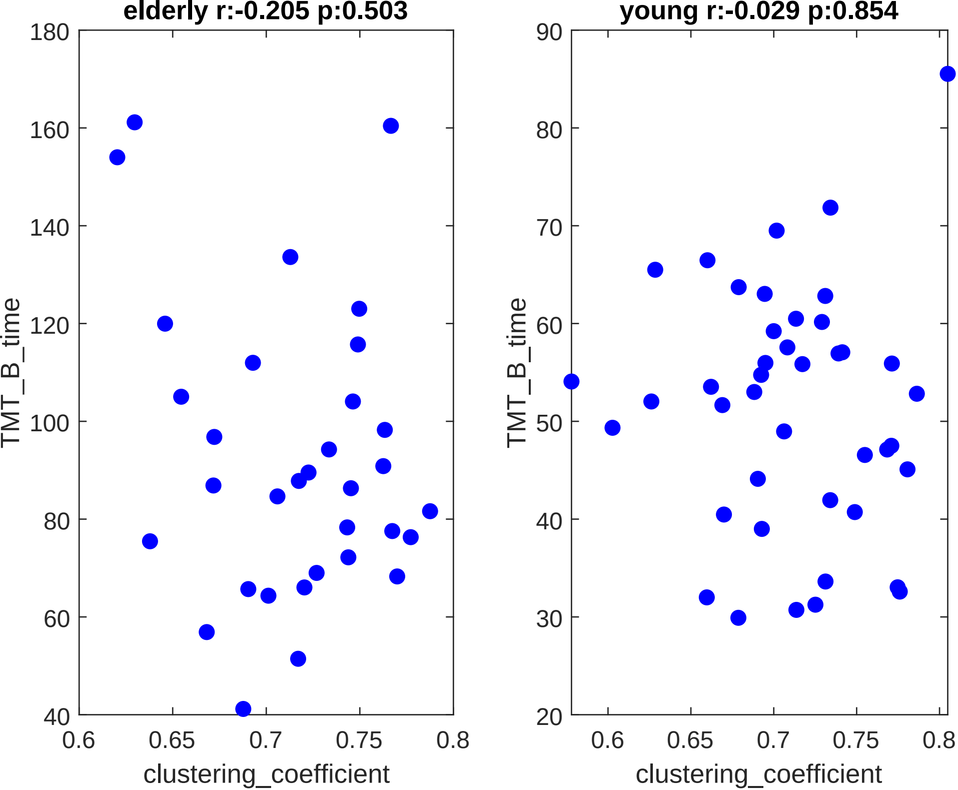

elderly r:-0.205 p:0.503
young r:-0.029 p:0.854
TMT_B_time
clustering_coefficient
180
160
140
120
100
80
60
40
90
70
50
30
20
0.6
0.65
0.7
0.75
0.8

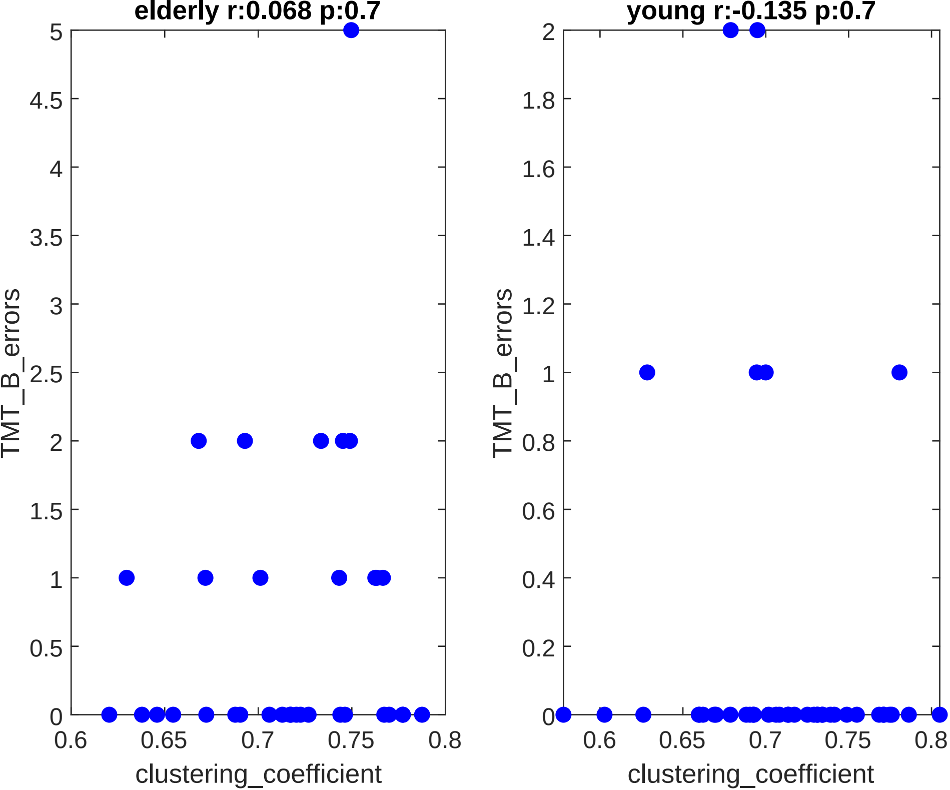

elderly r:0.068 p:0.7
TMT_B_errors
clustering_coefficient
0
0.5
1
1.5
2
2.5
3
3.5
4
4.5
5
0.6
0.65
0.7
0.75
0.8
young r:-0.135 p:0.7
TMT_B_errors
clustering_coefficient
0
0.2
0.4
0.6
0.8
1
1.2
1.4
1.6
1.8
2
0.6
0.65
0.7
0.75
0.8

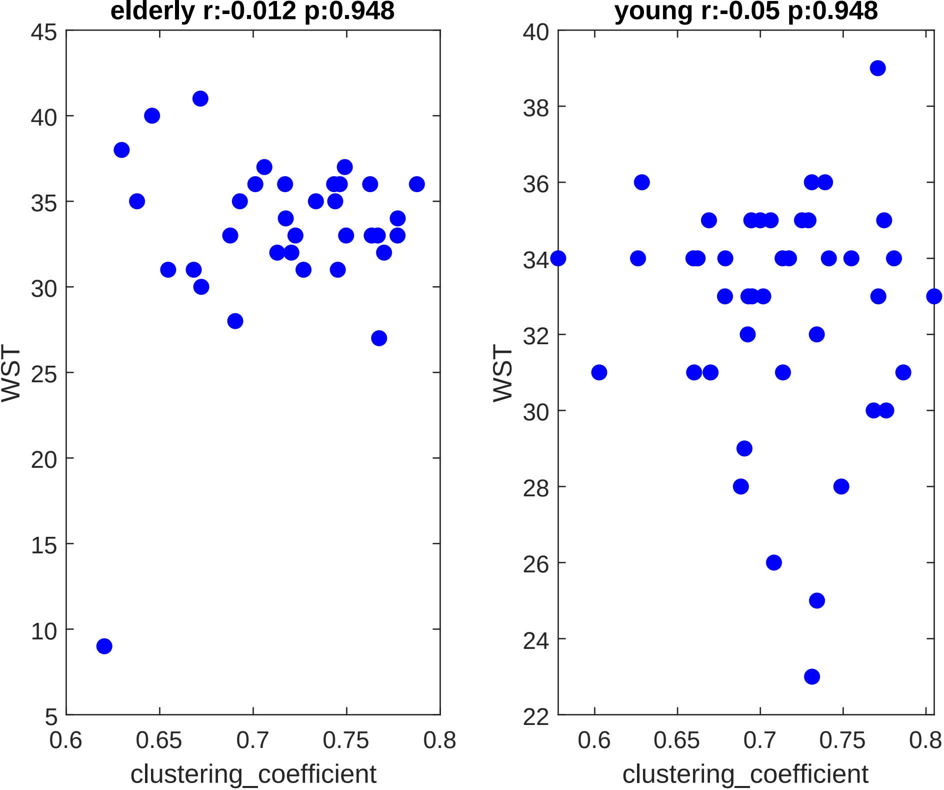
elderly r:-0.012 p:0.948
young r:-0.05 p:0.948
WST
clustering_coefficient
WST
clustering_coefficient
