## Supplementary material for "Whole-Body Networks: A Holistic Approach for Studying Aging": Correlations_node_degree.pdf

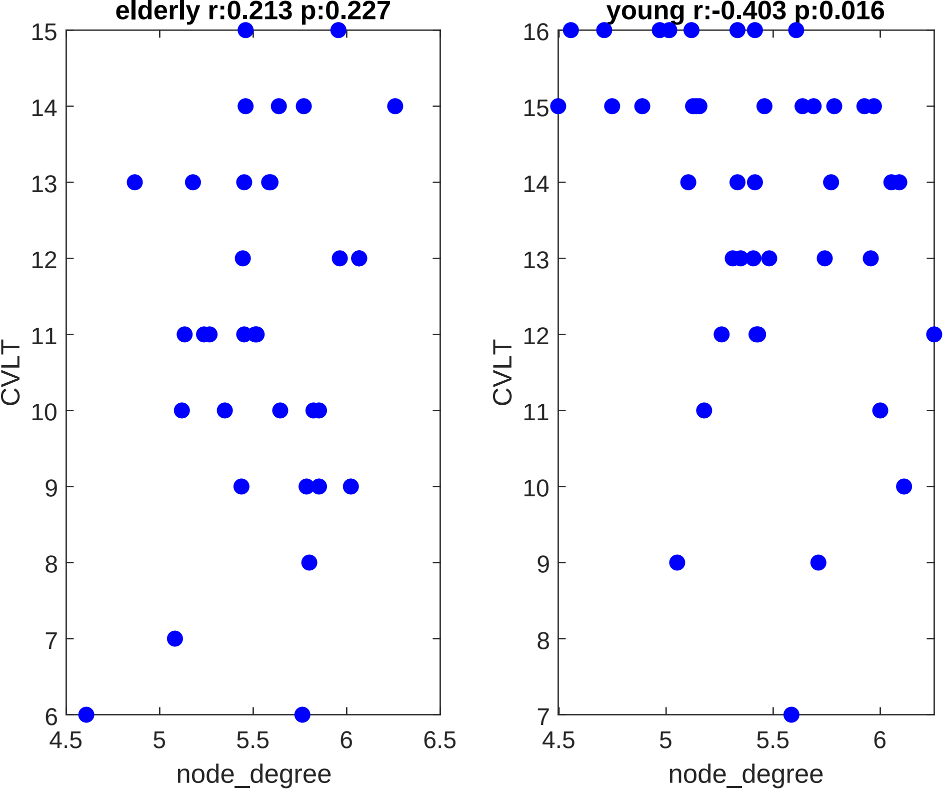

elderly r:0.213 p:0.227
young r:-0.403 p:0.016
CVLT
node_degree
CVLT
node_degree

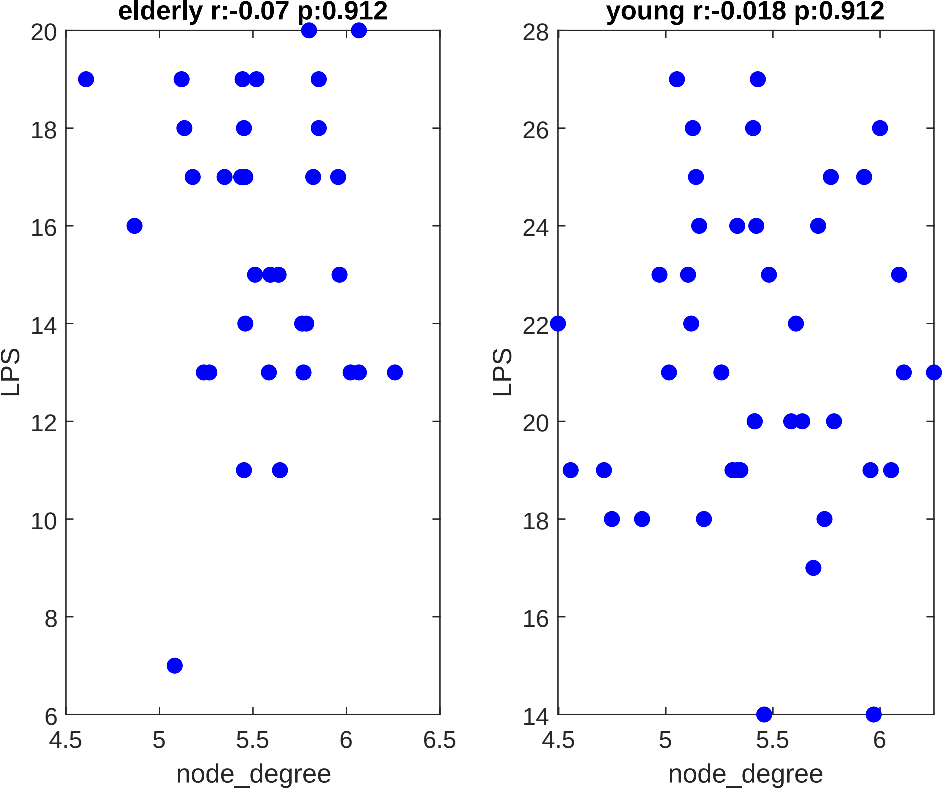
elderly r:-0.07 p:0.912
young r:-0.018 p:0.912
LPS
node_degree
20
18
16
14
12
10
8
6
4.5
5
5.5
6
6.5
28
26
24
22
20
18
16
14
4.5
5
5.5
6
LPS
node_degree

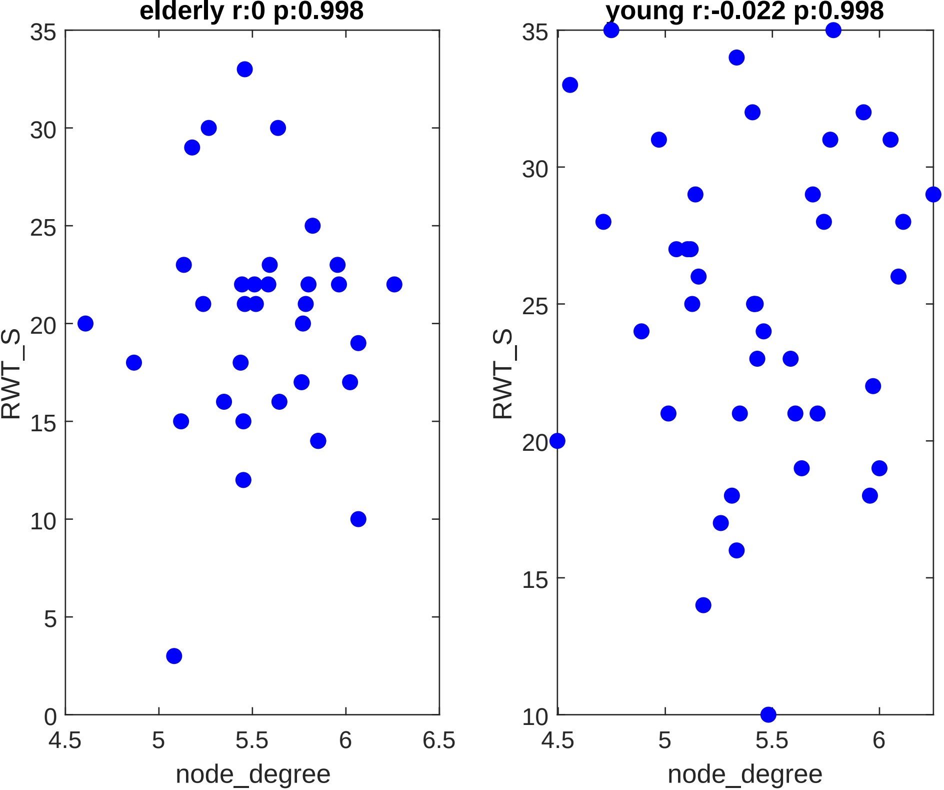

elderly r:0 p:0.998
young r:-0.022 p:0.998
RWT_S
node_degree
35
30
25
20
15
10
5
0
4.5
5
5.5
6
6.5

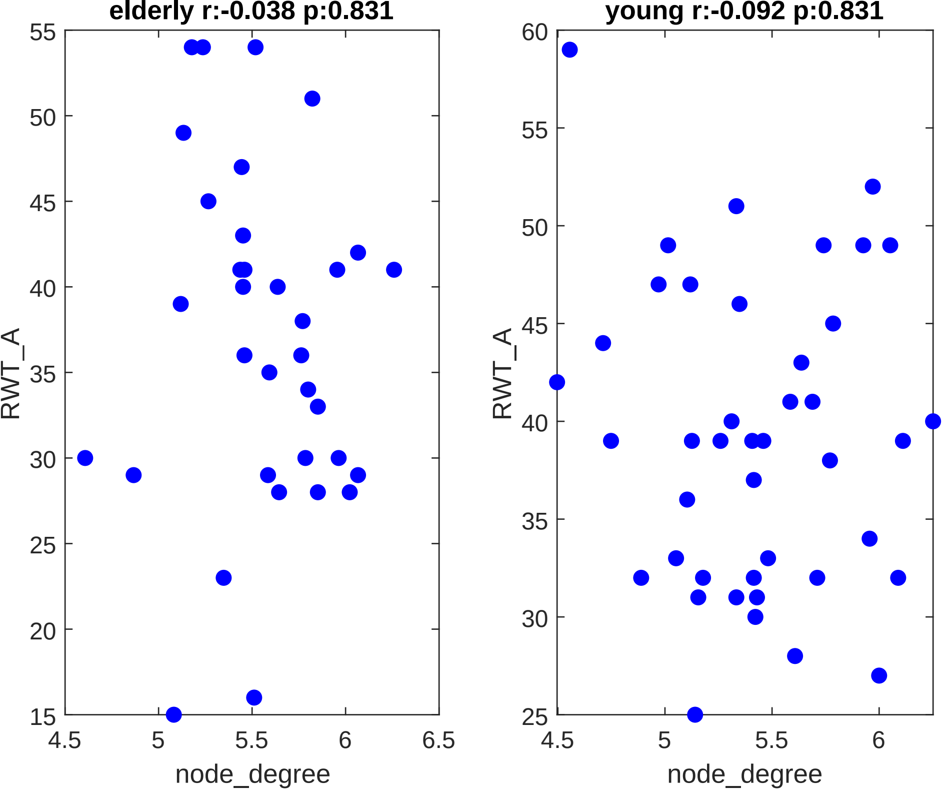

elderly r:-0.038 p:0.831
young r:-0.092 p:0.831
RWT_A
node_degree
55
50
45
40
35
30
25
20
15
60
4.5
5
5.5
6
6.5

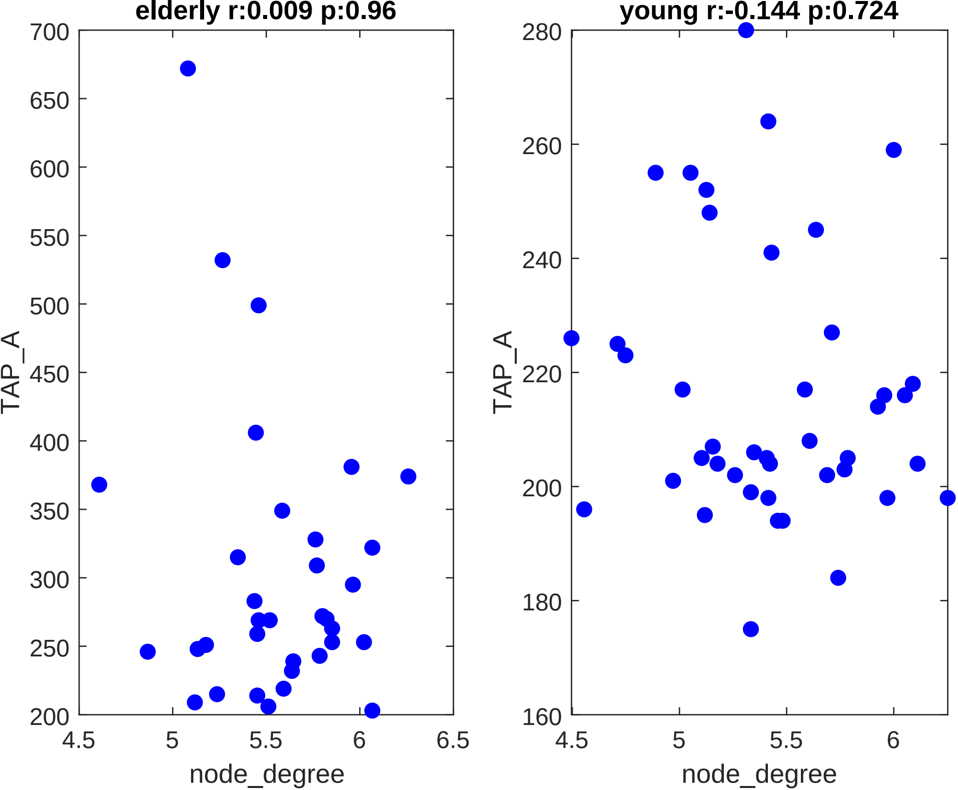
elderly r:0.009 p:0.96
young r:-0.144 p:0.724
TAP_A
node_degree
700
650
600
550
500
450
400
350
300
250
200
4.5
5
5.5
6
6.5
280
260
240
220
200
180
160

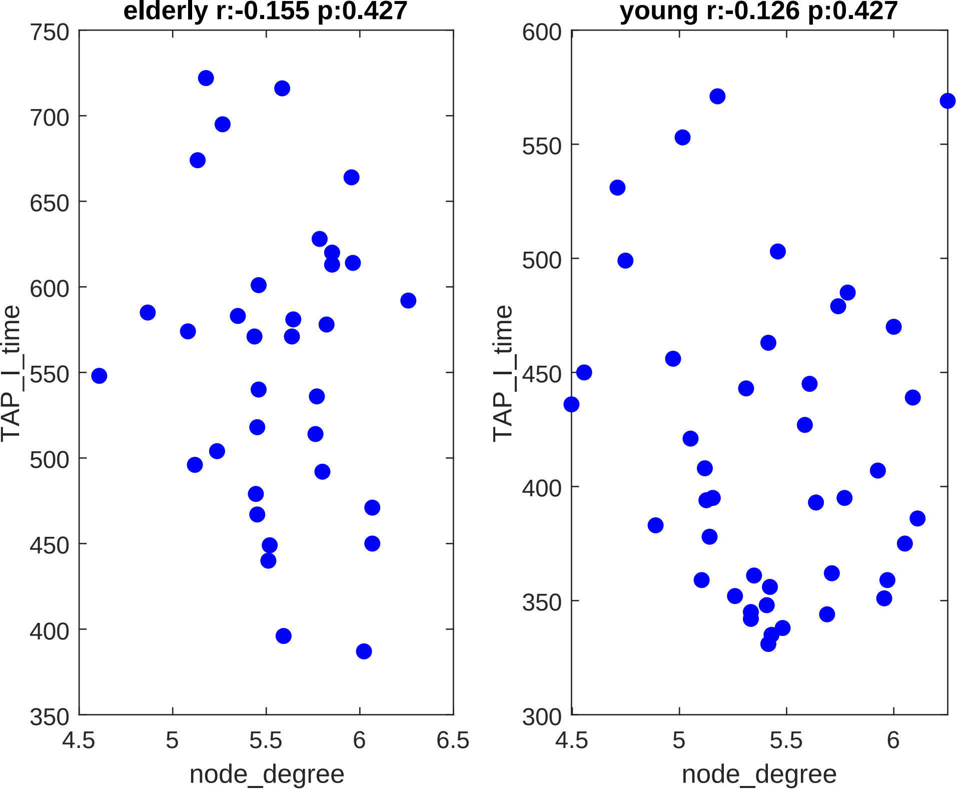
elderly r:-0.155 p:0.427
young r:-0.126 p:0.427
TAP_I_time
node_degree
750
700
650
600
550
500
450
400
350
300
4.5
5
5.5
6
6.5

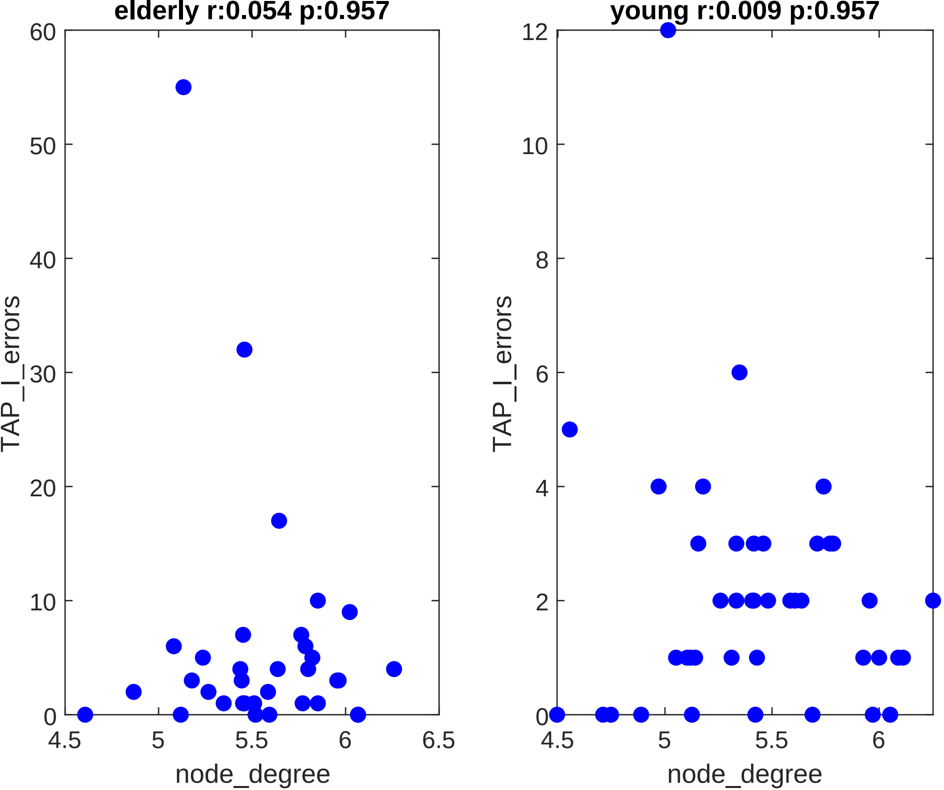
elderly r:0.054 p:0.957
young r:0.009 p:0.957
TAP_I_errors
node_degree
TAP_I_errors
node_degree
0
10
20
30
40
50
60
4.5
5
5.5
6
6.5
0
2
4
6
8
10
12
4.5
5
5.5
6

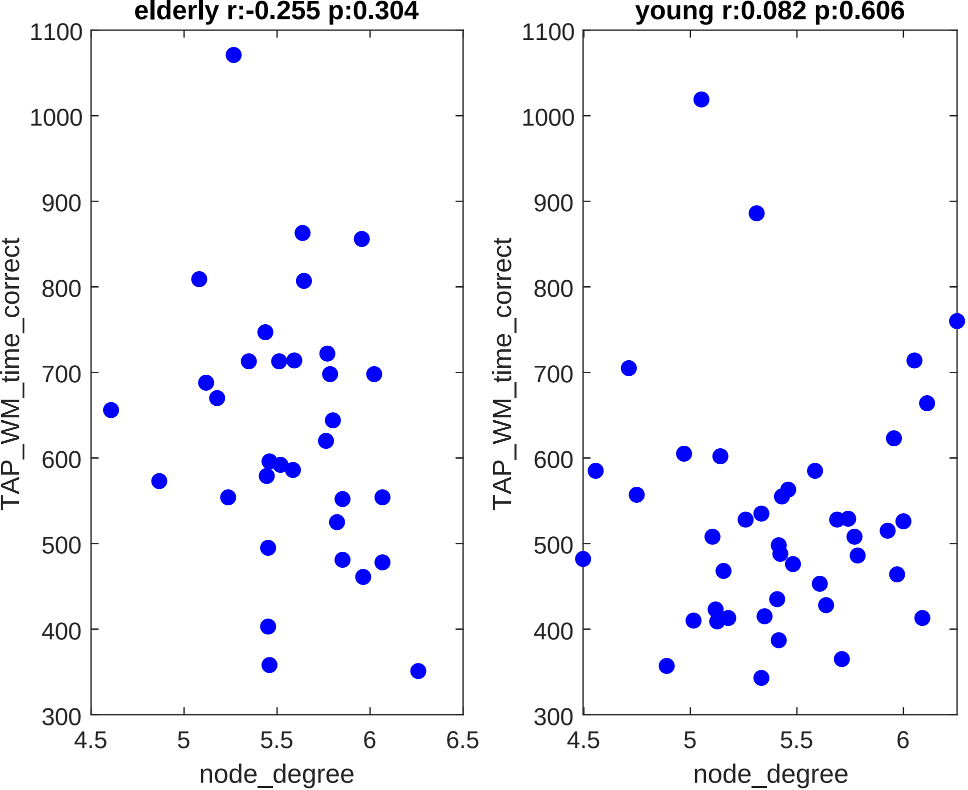

elderly r:-0.255 p:0.304
young r:0.082 p:0.606
1100
1000
900
800
700
600
500
400
300
4.5
5
5.5
6
6.5
TAP_WM_time_correct
node_degree

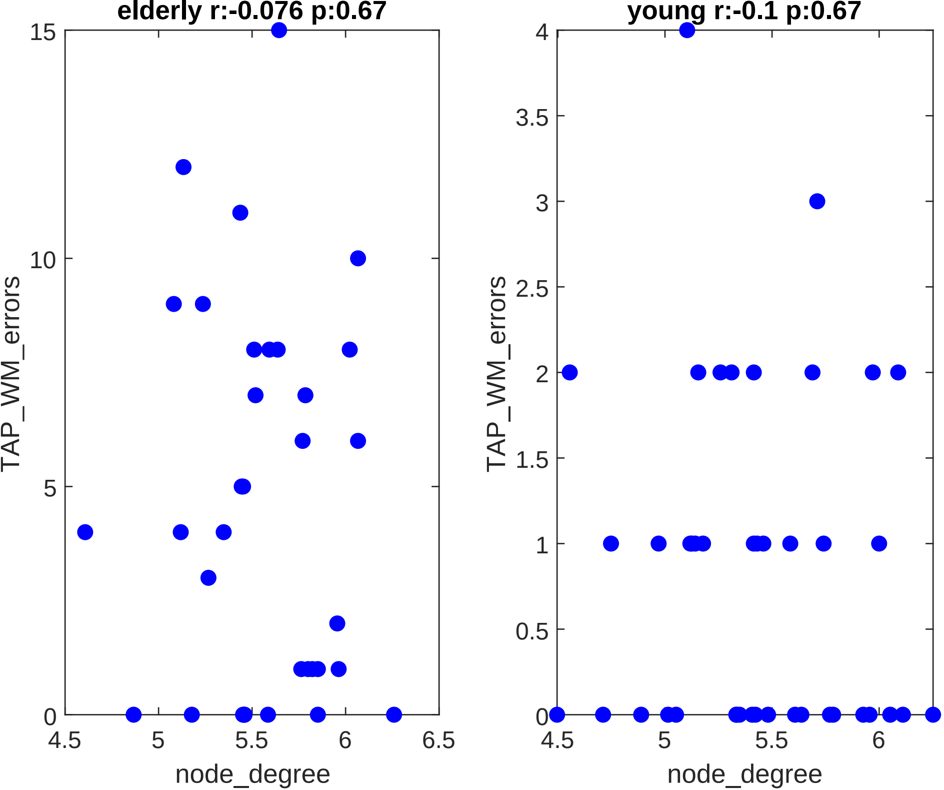
elderly r:-0.076 p:0.67
young r:-0.1 p:0.67
TAP_WM_errors
TAP_WM_errors
node_degree
node_degree
15
10
5
0
4
3.5
3
2.5
2
1.5
1
0.5
0
4.5
5
5.5
6
6.5
4.5
5
5.5
6

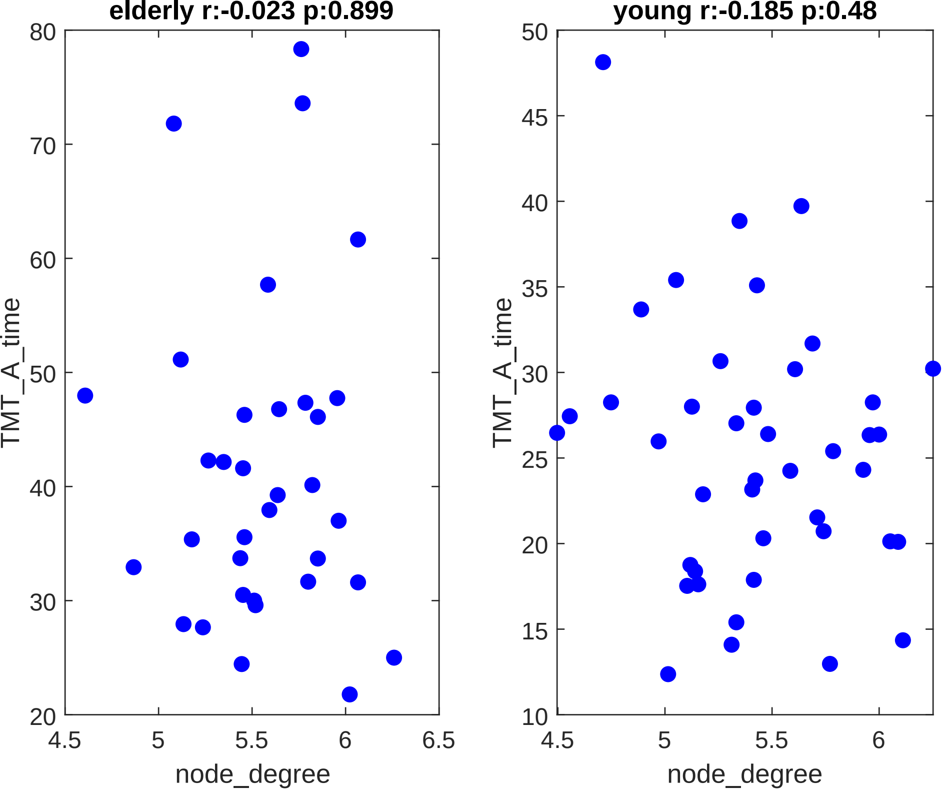
elderly r:-0.023 p:0.899
young r:-0.185 p:0.48
TMT_A_time
node_degree
80
70
60
50
40
30
20
4.5
5
5.5
6
6.5
TMT_A_time
node_degree
50
45
40
35
30
25
20
15
10
4.5
5
5.5
6

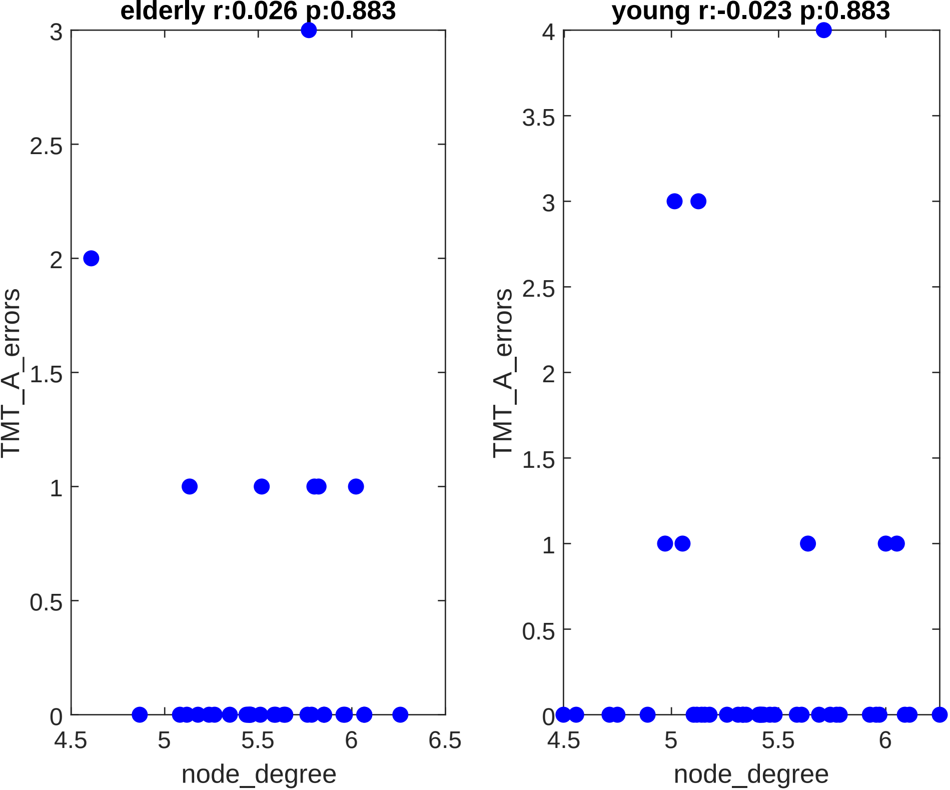
elderly r:0.026 p:0.883
TMT_A_errors
node_degree
3
2.5
2
1.5
1
0.5
0
4.5
5
5.5
6
6.5
young r:-0.023 p:0.883
TMT_A_errors
node_degree
4
3.5
3
2.5
2
1.5
1
0.5
0
4.5
5
5.5
6

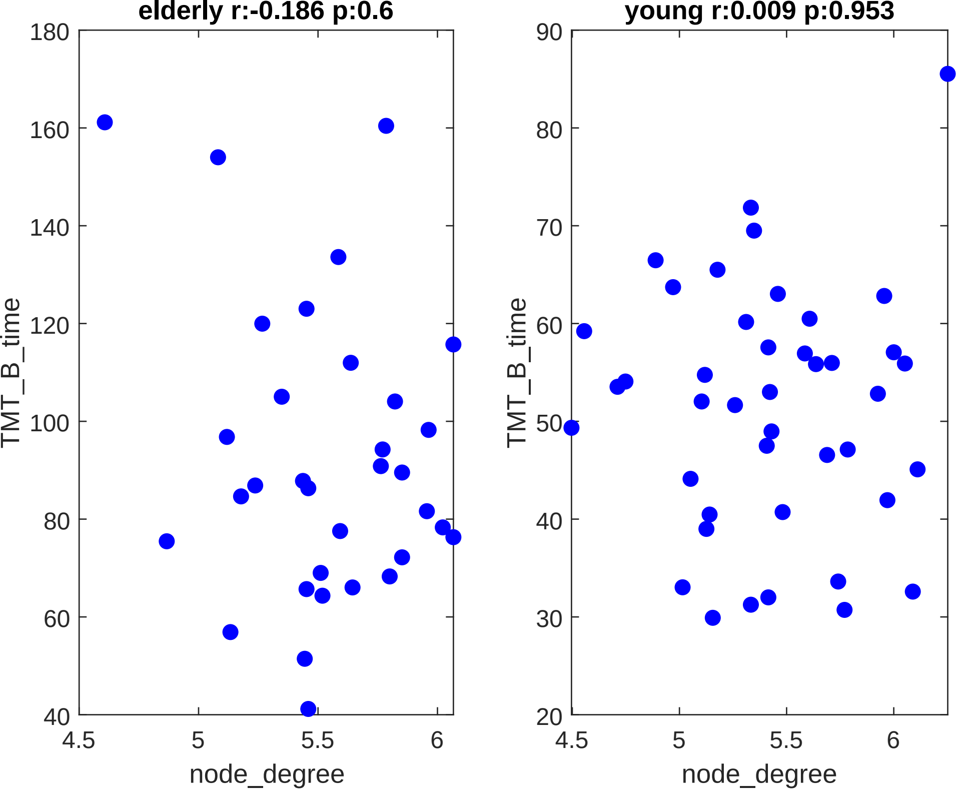
elderly r:-0.186 p:0.6
young r:0.009 p:0.953
TMT_B_time
node_degree
180
160
140
120
100
80
60
40
90
70
50
30
20
4.5
5
5.5
6

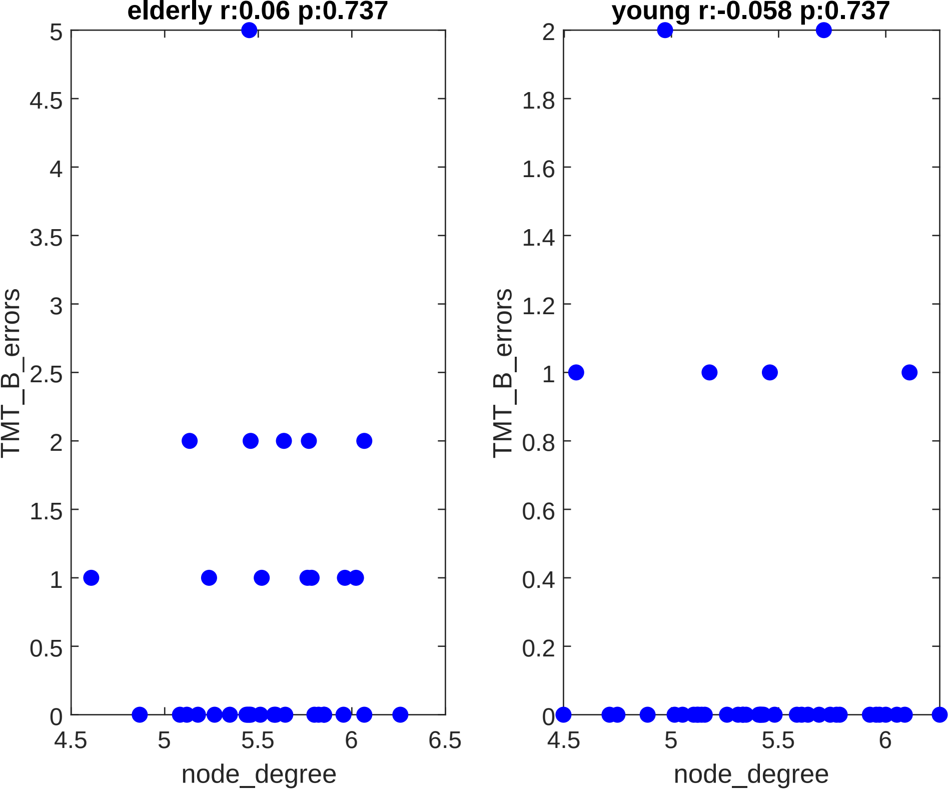
elderly r:0.06 p:0.737
young r:-0.058 p:0.737
TMT_B_errors
node_degree
0
0.5
1
1.5
2
2.5
3
3.5
4
4.5
5
4.5
5
5.5
6
6.5
0
0.2
0.4
0.6
0.8
1
1.2
1.4
1.6
1.8
2
4.5
5
5.5
6

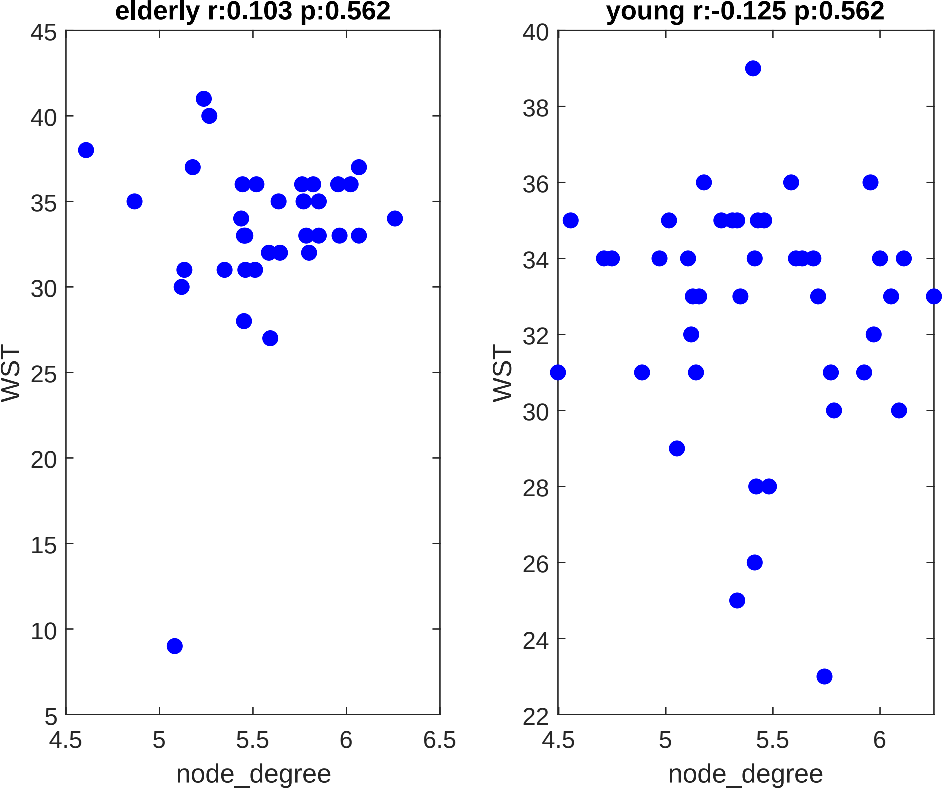

elderly r:0.103 p:0.562
45
40
35
30
25
20
15
10
5
4.5
5
5.5
6
6.5
WST
node_degree
young r:-0.125 p:0.562
40
38
36
34
32
30
28
26
24
22
4.5
5
5.5
6
WST
node_degree
