## Supplementary material for "Whole-Body Networks: A Holistic Approach for Studying Aging": Correlations_path_length.pdf

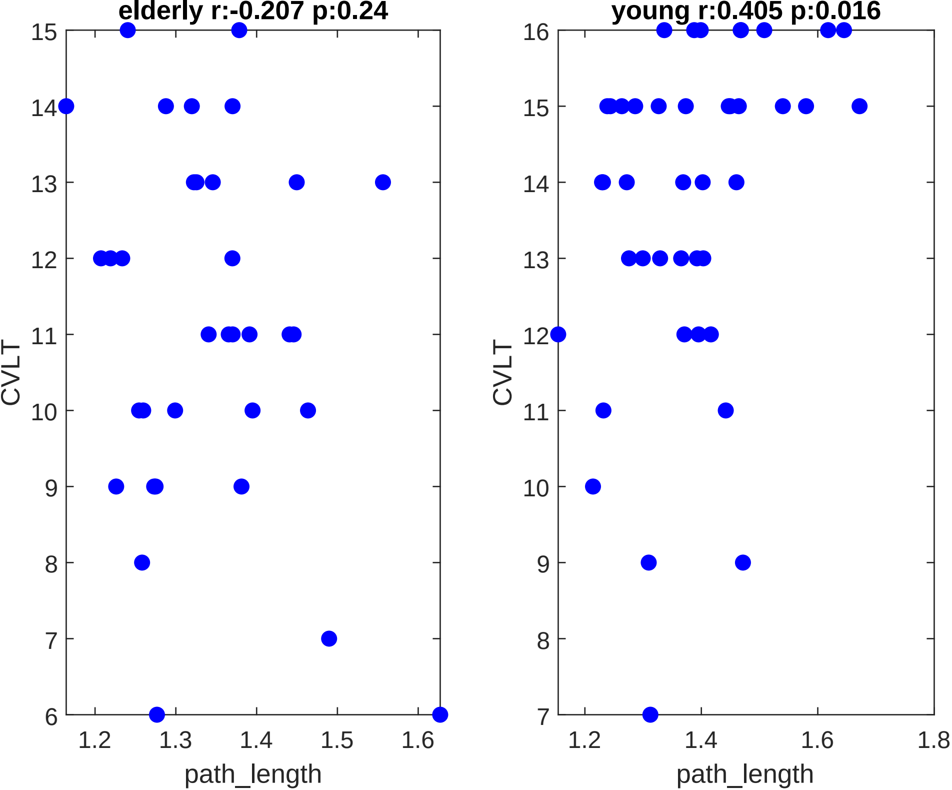
elderly r:-0.207 p:0.24
young r:0.405 p:0.016
CVLT
CVLT
path_length
path_length
15
14
13
12
11
10
9
8
7
6
16
15
14
13
12
11
10
9
8
7
1.2
1.3
1.4
1.5
1.6
1.2
1.4
1.6
1.8

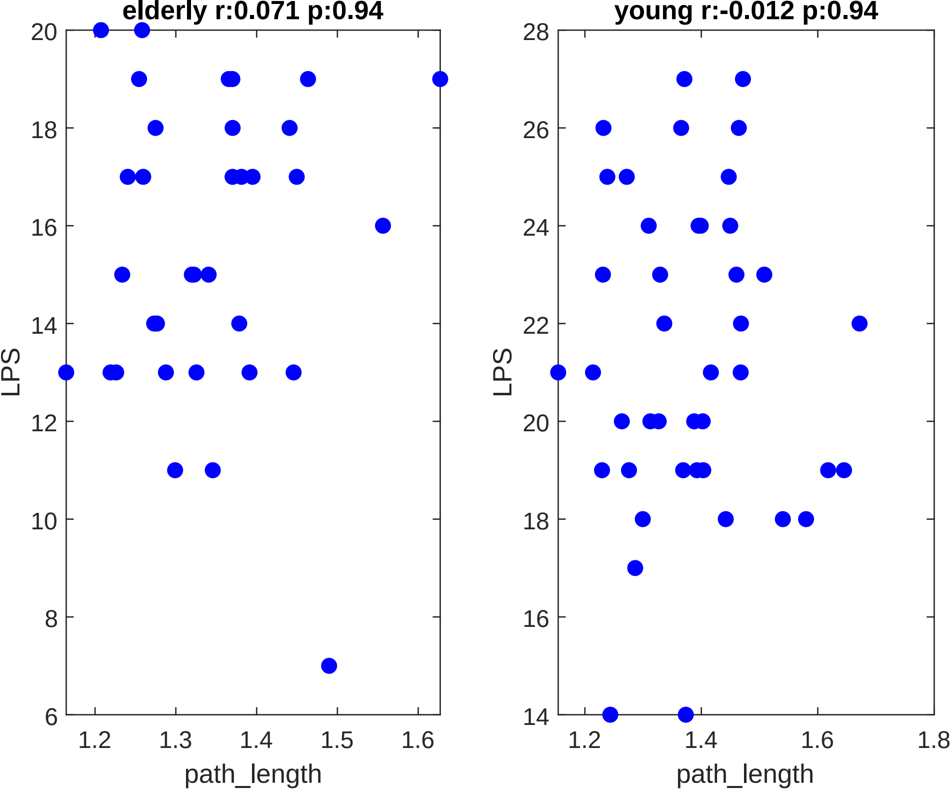

elderly r:0.071 p:0.94
young r:-0.012 p:0.94
LPS
path_length
20
18
16
14
12
10
8
6
1.2
1.3
1.4
1.5
1.6
28
26
24
22
20
18
16
14
1.2
1.4
1.6
1.8

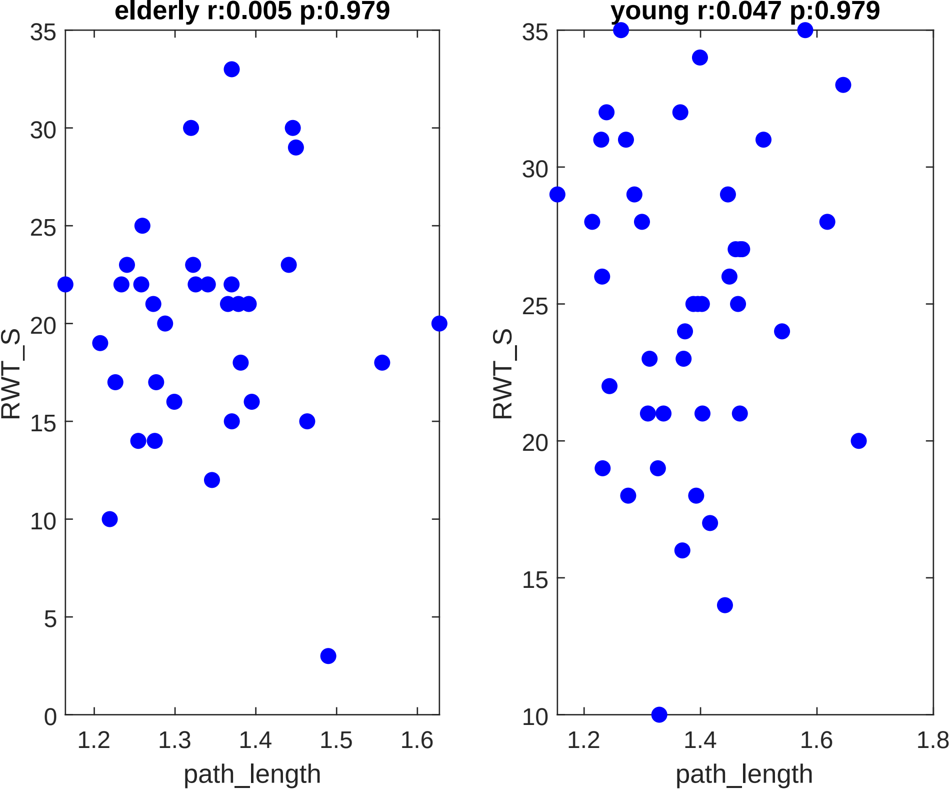

elderly r:0.005 p:0.979
young r:0.047 p:0.979
RWT_S
path_length
RWT_S
path_length

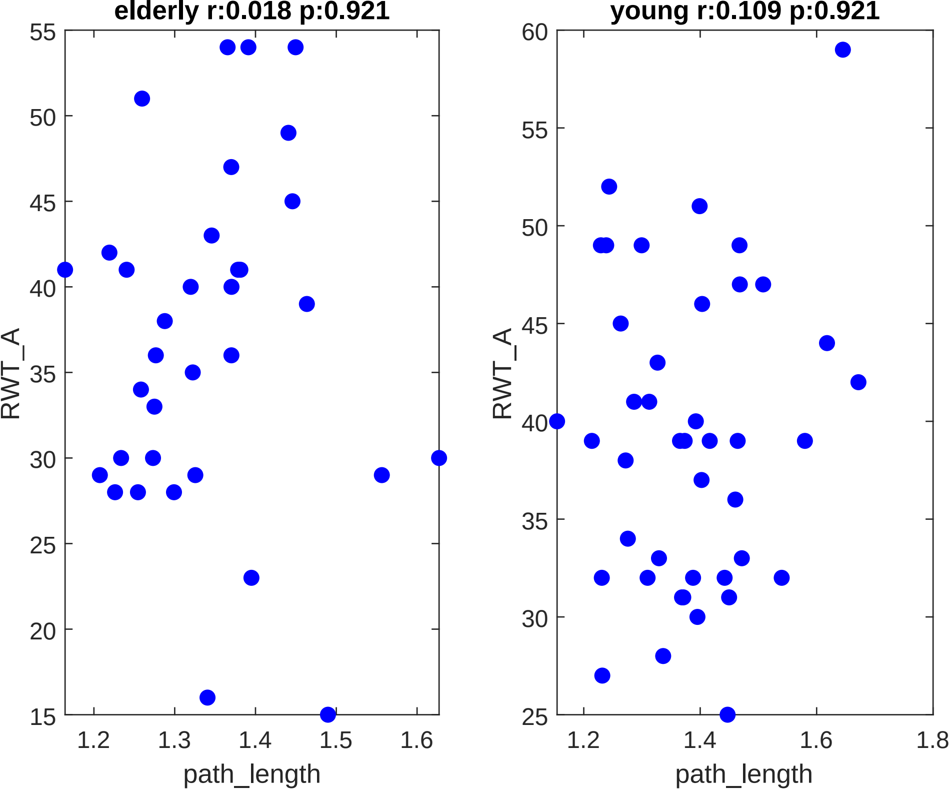
elderly r:0.018 p:0.921
RWT_A
path_length
55
50
45
40
35
30
25
20
15
1.2
1.3
1.4
1.5
1.6
young r:0.109 p:0.921
RWT_A
path_length
60
55
50
45
40
35
30
25
1.2
1.4
1.6
1.8

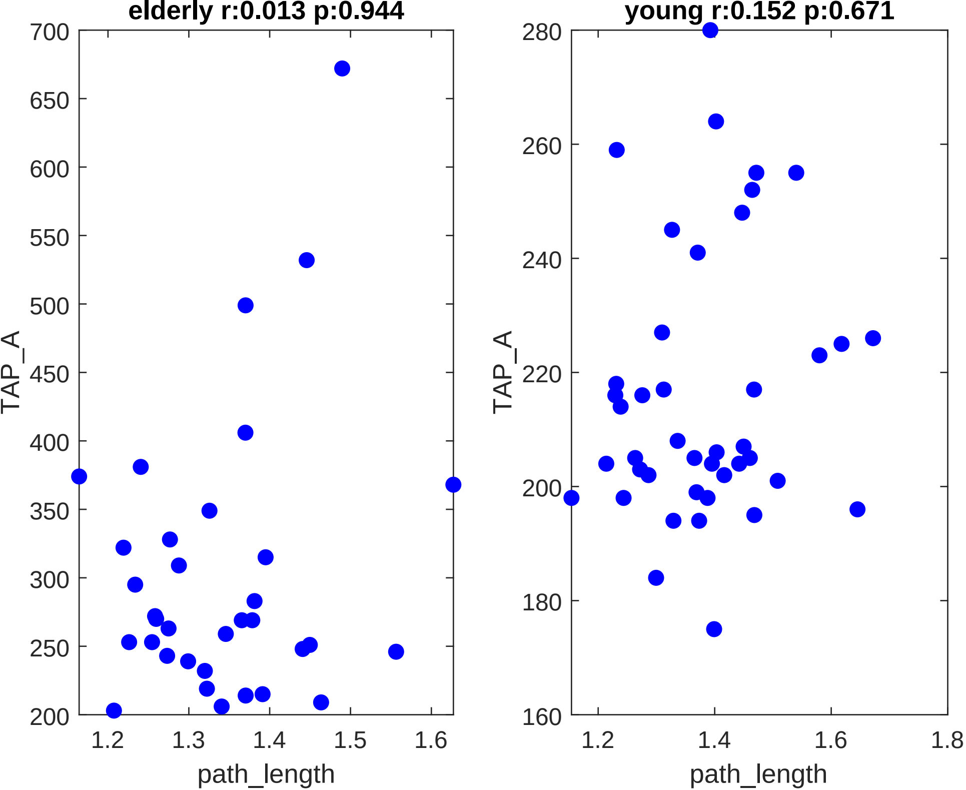

elderly r:0.013 p:0.944
young r:0.152 p:0.671
TAP_A
path_length
700
650
600
550
500
450
400
350
300
250
200
1.2
1.3
1.4
1.5
1.6
280
260
240
220
200
180
160
1.8

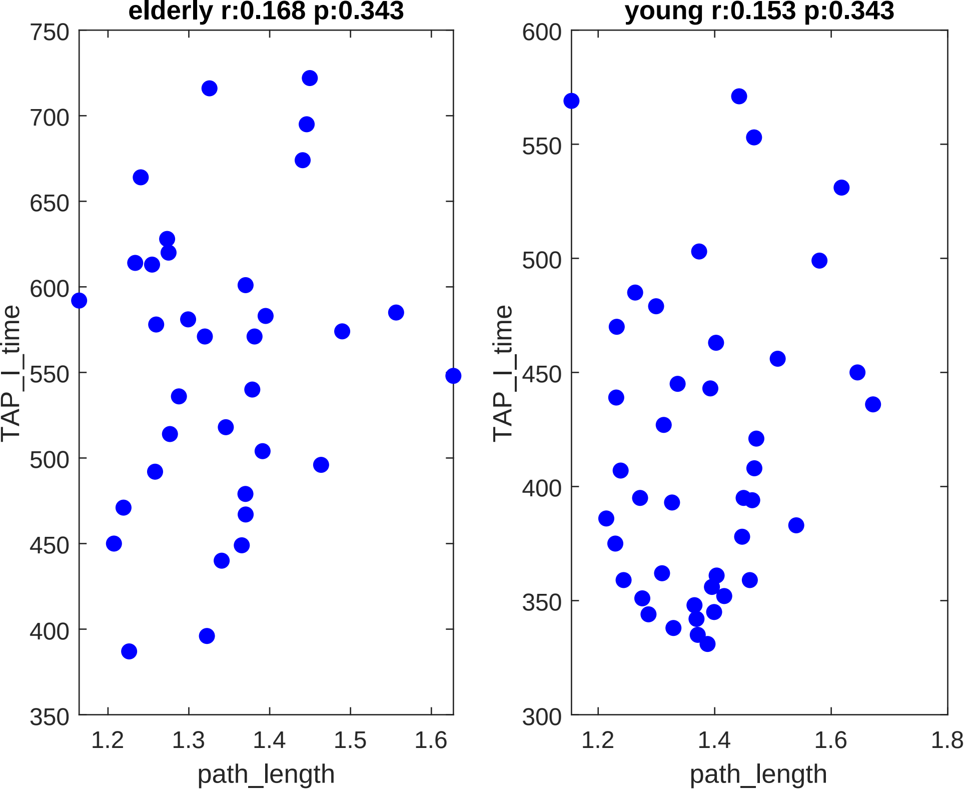

elderly r:0.168 p:0.343
young r:0.153 p:0.343
TAP_I_time
path_length
750
700
650
600
550
500
450
400
350
300
1.2
1.3
1.4
1.5
1.6
1.8

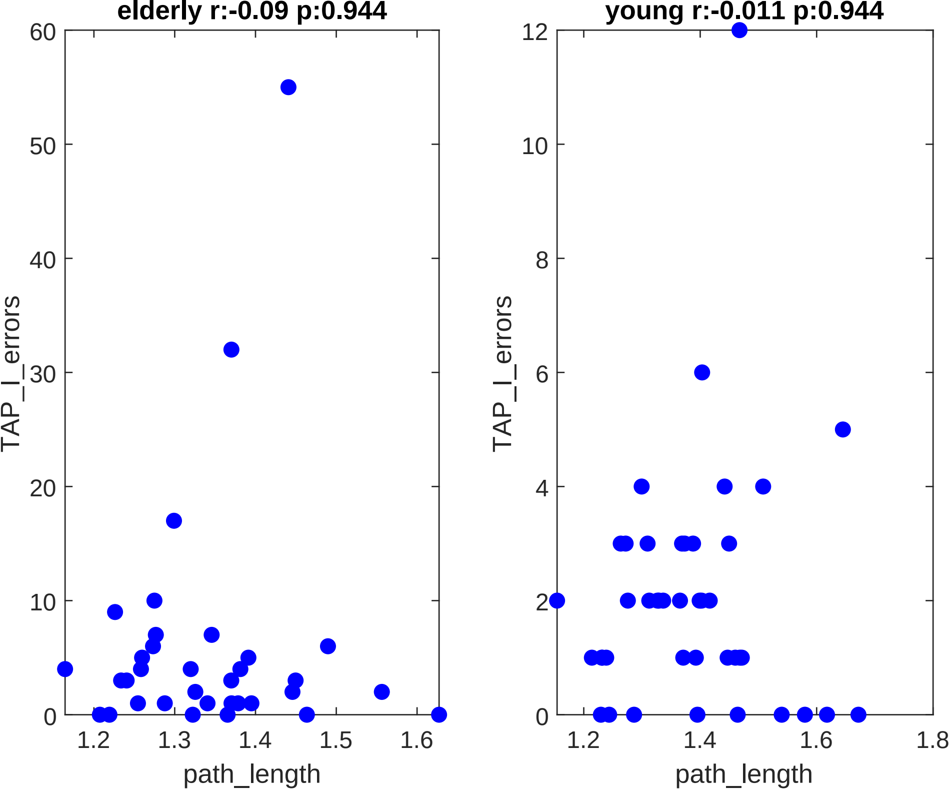

elderly r:-0.09 p:0.944
TAP_I_errors
path_length
60
50
40
30
20
10
0
1.2
1.3
1.4
1.5
1.6
young r:-0.011 p:0.944
TAP_I_errors
path_length
12
10
8
6
4
2
0
1.2
1.4
1.6
1.8

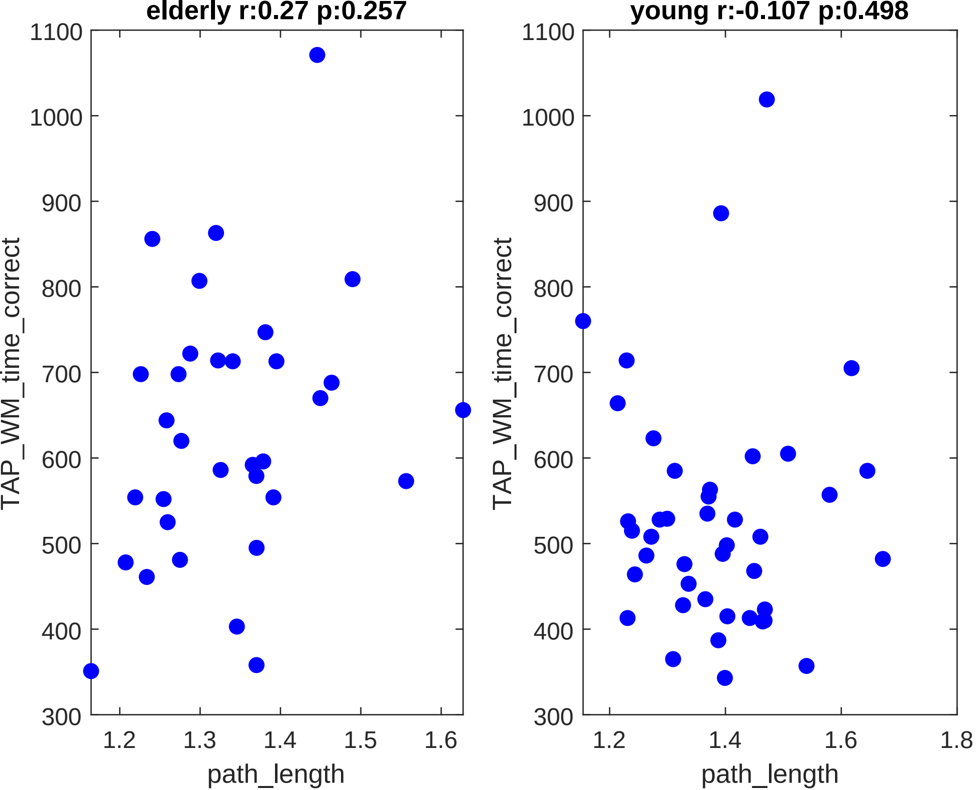

elderly r:0.27 p:0.257
young r:-0.107 p:0.498
TAP_WM_time_correct
path_length
1100
1000
900
800
700
600
500
400
300
1.2
1.3
1.4
1.5
1.6
1.8

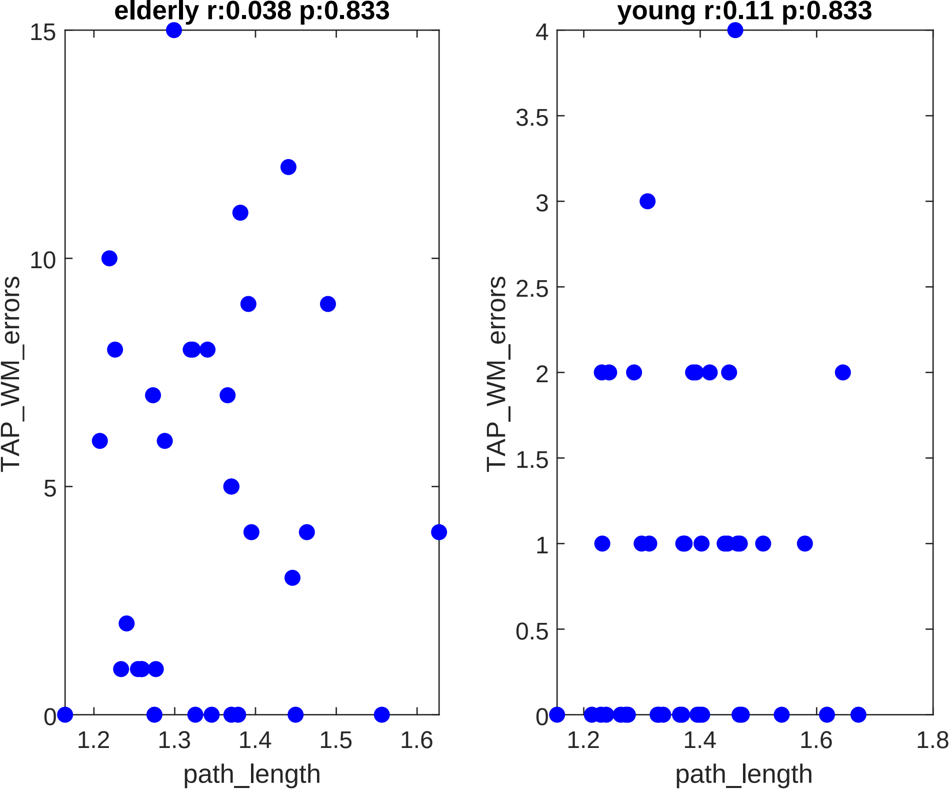

elderly r:0.038 p:0.833
young r:0.11 p:0.833
TAP_WM_errors
path_length
15
10
5
0
1.2
1.3
1.4
1.5
1.6
4
3.5
3
2.5
2
1.5
1
0.5
0
1.2
1.4
1.6
1.8

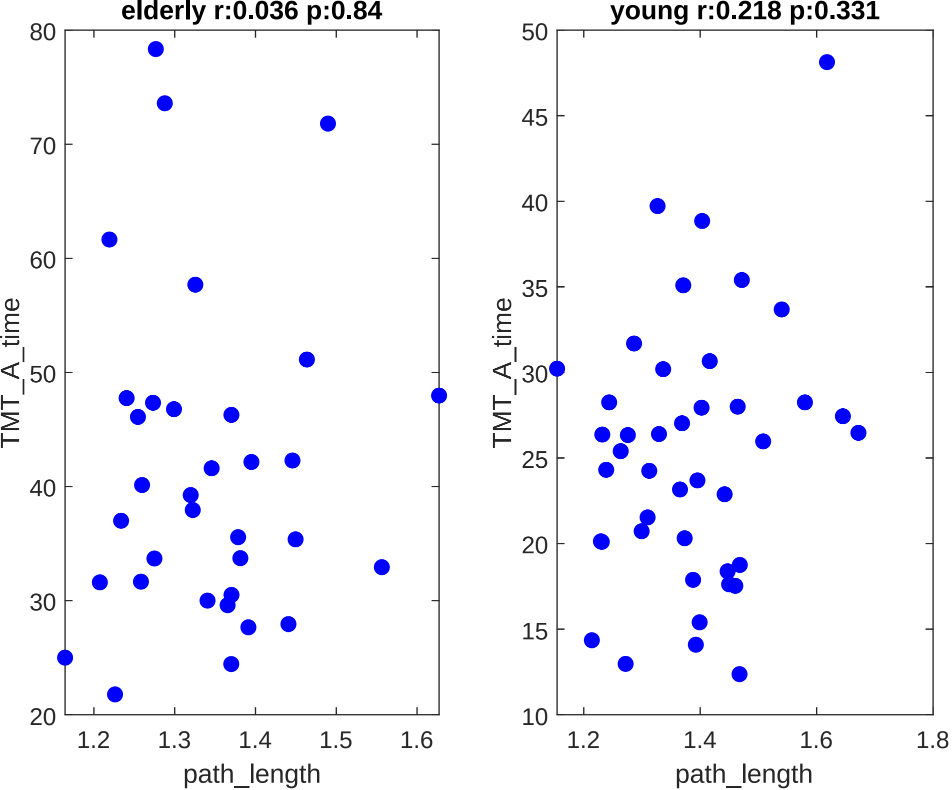

elderly r:0.036 p:0.84
young r:0.218 p:0.331
TMT_A_time
path_length
20
30
40
50
60
70
80
1.2
1.3
1.4
1.5
1.6
10
15
20
25
30
35
40
45
50
1.2
1.4
1.6
1.8

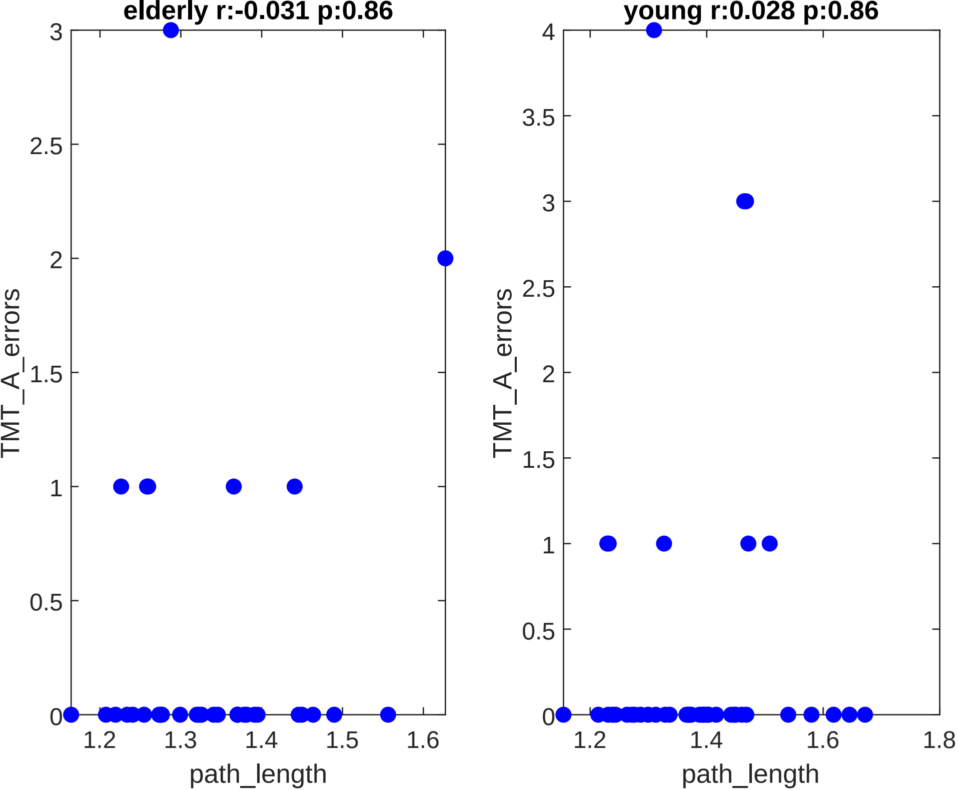
elderly r:-0.031 p:0.86
young r:0.028 p:0.86
TMT_A_errors
path_length
0
0.5
1
1.5
2
2.5
3
3.5
4
1.2
1.3
1.4
1.5
1.6
1.8

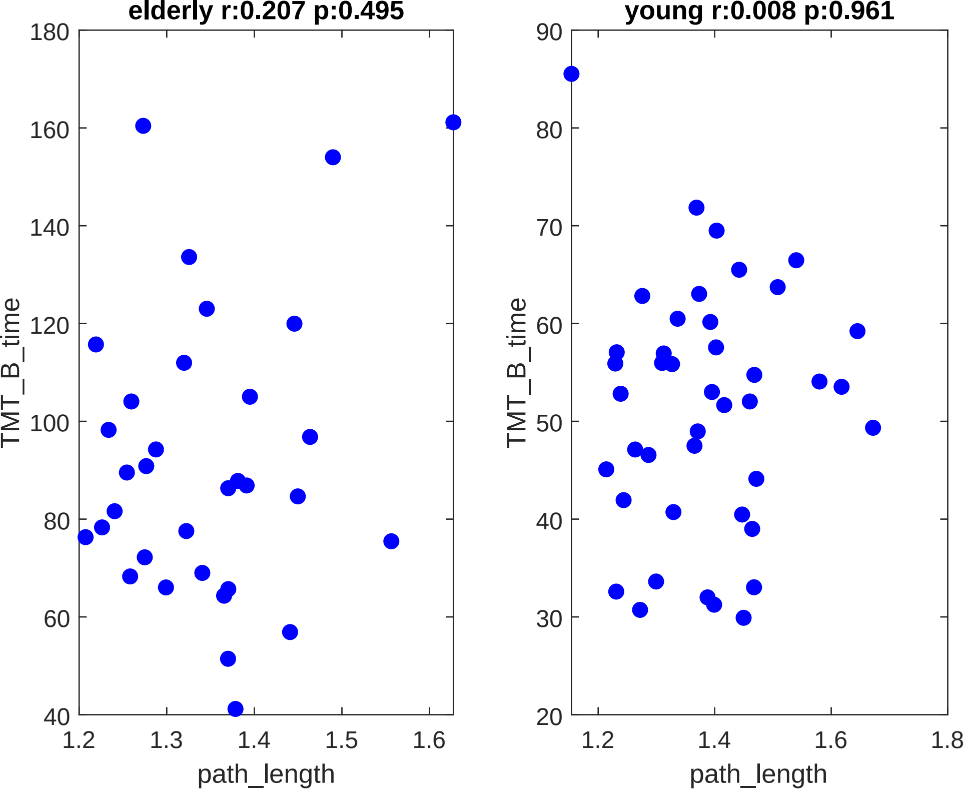
elderly r:0.207 p:0.495
young r:0.008 p:0.961
TMT_B_time
path_length
180
160
140
120
100
80
60
40
1.2
1.3
1.4
1.5
1.6
90
70
50
30
20
1.8

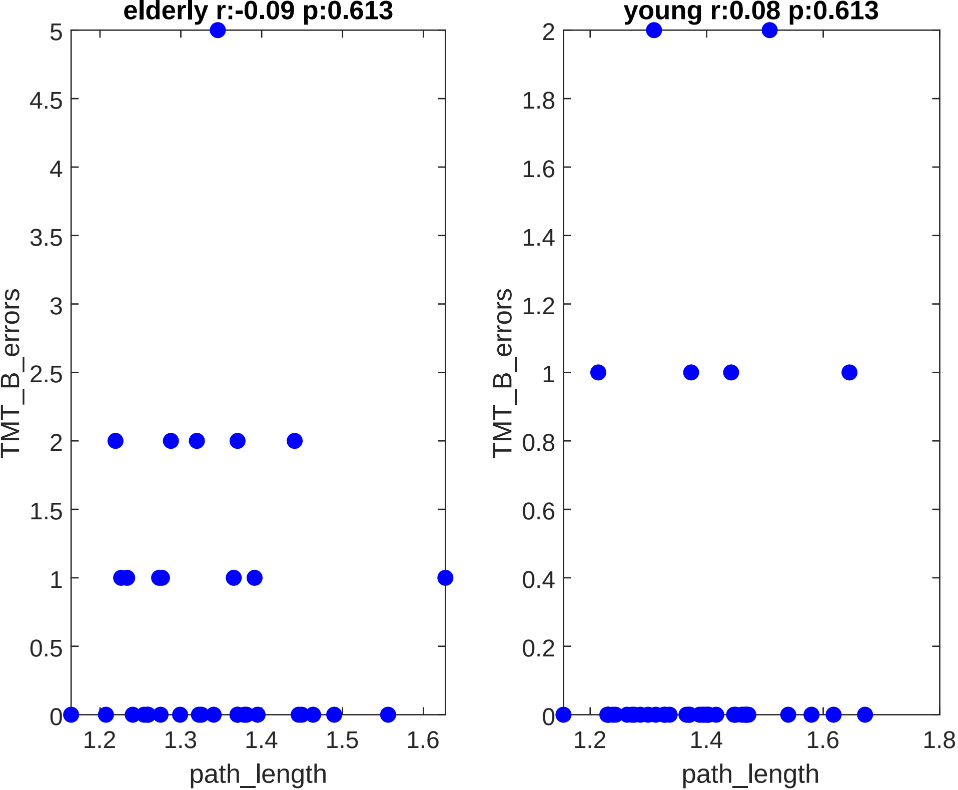

elderly r:-0.09 p:0.613
young r:0.08 p:0.613
TMT_B_errors
path_length
5
4.5
4
3.5
3
2.5
2
1.5
1
0.5
0
1.2
1.3
1.4
1.5
1.6
1.8
0.8
0.6
0.4
0.2

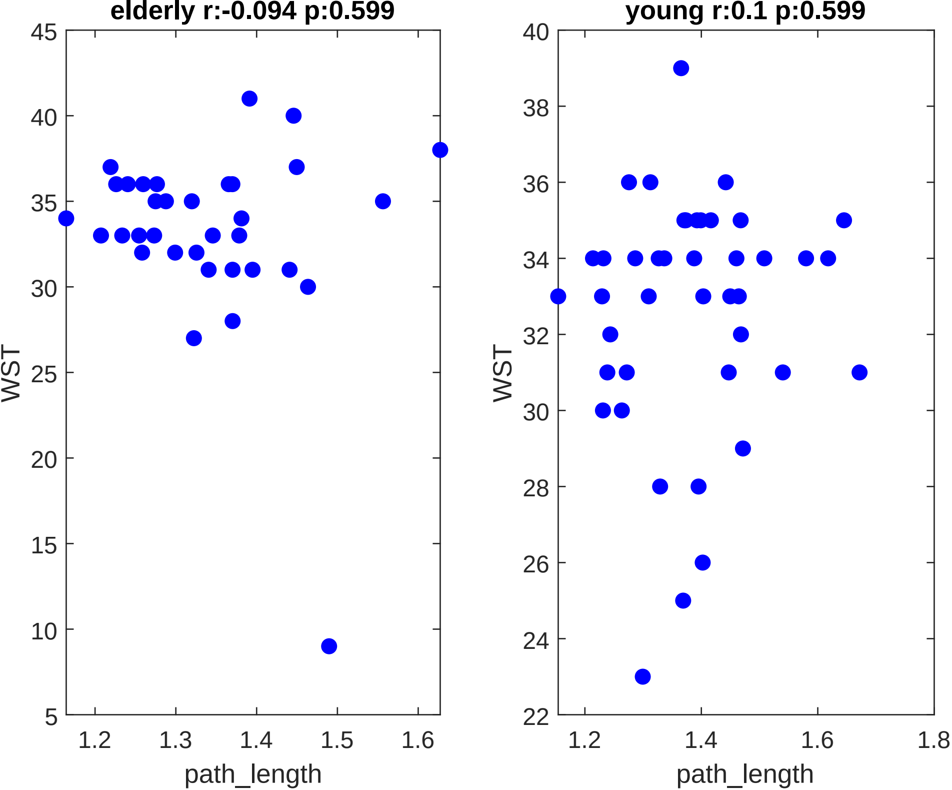
elderly r:-0.094 p:0.599
young r:0.1 p:0.599
WST
path_length
45
40
35
30
25
20
15
10
5
1.2
1.3
1.4
1.5
1.6
38
36
34
32
28
26
24
22
1.8
