## Supplementary figures and images for "Whole-Body Networks: A Holistic Approach for Studying Aging"

### Whole-Body Networks: A Holistic Approach for Studying Aging

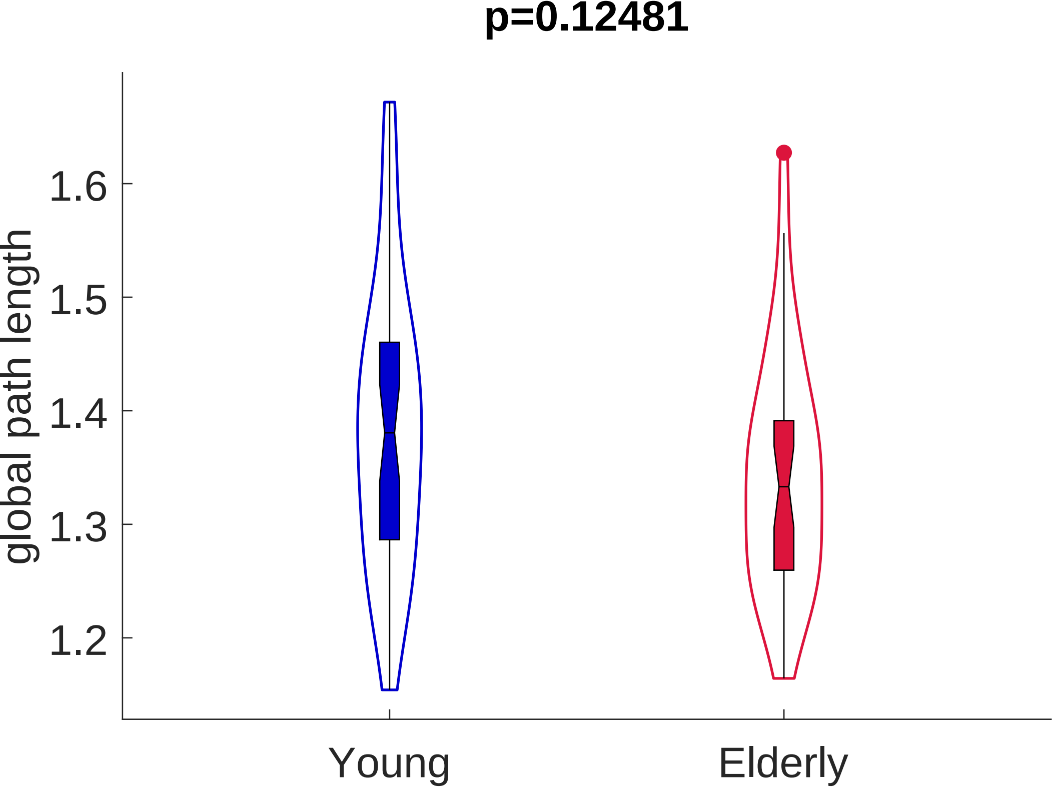

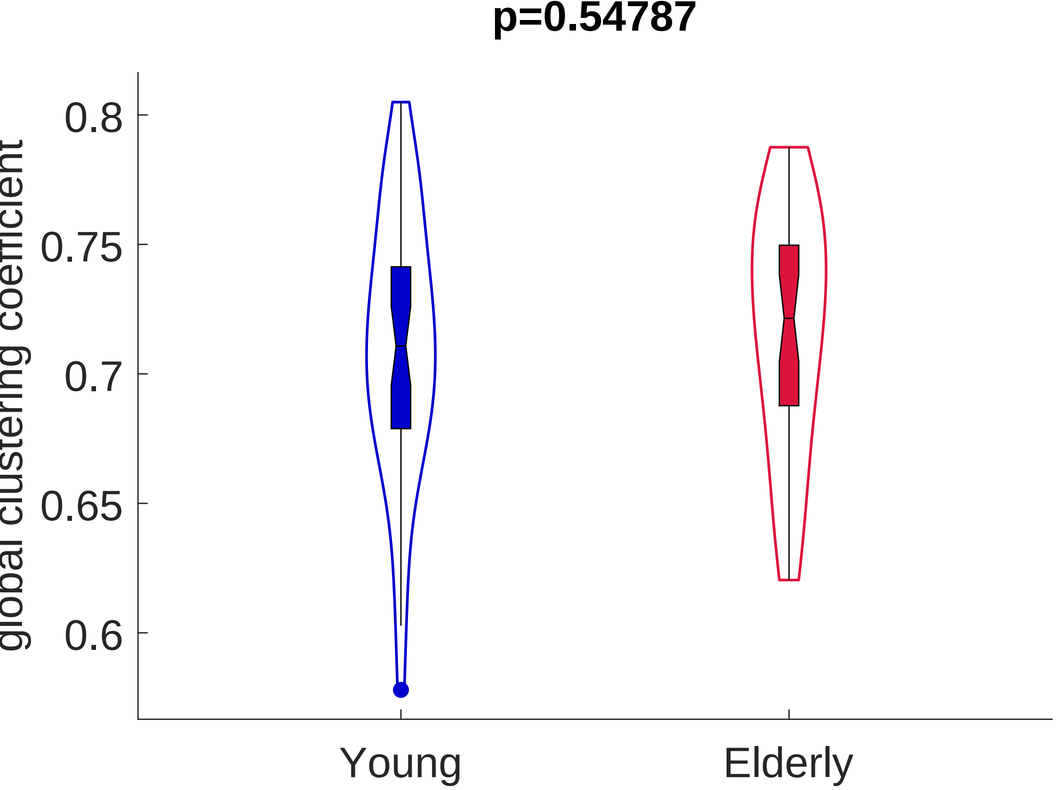

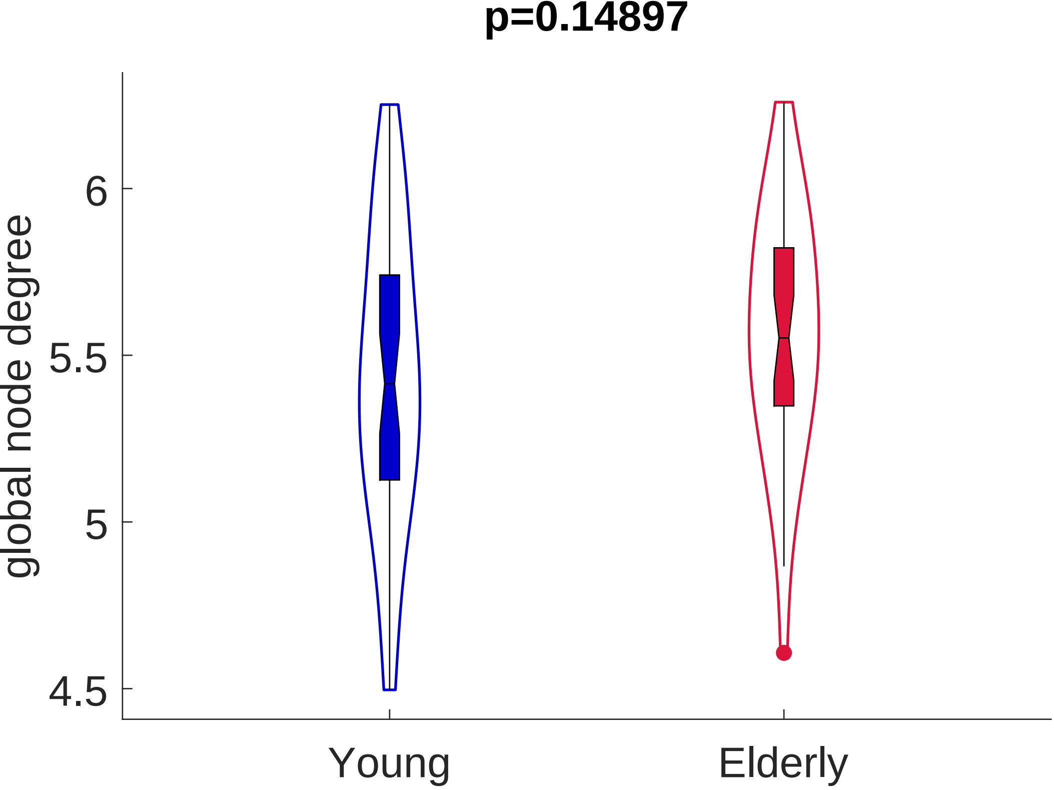
